# RF coil design strategies for improving SNR at the ultrahigh magnetic field of 10.5 Tesla

**DOI:** 10.1101/2024.05.23.595628

**Authors:** Matt Waks, Russell L. Lagore, Edward Auerbach, Andrea Grant, Alireza Sadeghi-Tarakameh, Lance DelaBarre, Steve Jungst, Nader Tavaf, Riccardo Lattanzi, Ilias Giannakopoulos, Steen Moeller, Xiaoping Wu, Essa Yacoub, Luca Vizioli, Simon Schmidt, Gregory J. Metzger, Yigitcan Eryaman, Gregor Adriany, Kamil Uğurbil

## Abstract

**Purpose:** To develop multichannel transmit and receive arrays towards capturing the ultimate-intrinsic-SNR (uiSNR) at 10.5 Tesla (T) and to demonstrate the feasibility and potential of whole-brain, high-resolution human brain imaging at this high field strength.

**Methods:** A dual row 16-channel self-decoupled transmit (Tx) array was converted to a 16Tx/Rx transceiver using custom transmit/receive switches. A 64-channel receive-only (64Rx) array was built to fit into the 16Tx/Rx array. Electromagnetic modeling and experiments were employed to define safe operation limits of the resulting 16Tx/80Rx array and obtain FDA approval for human use.

**Results:** The 64Rx array alone captured approximately 50% of the central uiSNR at 10.5T while the identical 7T 64Rx array captured ∼76% of uiSNR at this lower field strength. The 16Tx/80Rx configuration brought the fraction of uiSNR captured at 10.5T to levels comparable to the performance of the 64Rx array at 7T. SNR data obtained at the two field strengths with these arrays displayed 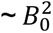 dependent increases over a large central region. Whole-brain high resolution T_2_* and T_1_ weighted anatomical and gradient-recalled echo EPI BOLD fMRI images were obtained at 10.5T for the first time with such an advanced array, illustrating the promise of >10T fields in studying the human brain.

**Conclusion:** We demonstrated the ability to approach the uiSNR at 10.5T over the human brain with a novel, high channel count array, achieving large SNR gains over 7T, currently the most commonly employed ultrahigh field platform, and demonstrate high resolution and high contrast anatomical and functional imaging at 10.5T.

## 1. INTRODUCTION

The introduction of ultrahigh magnetic fields for human applications was motivated by anticipated improvements in sensitivity, accuracy, and spatial resolution of functional brain imaging (fMRI) (e.g., reviews (1–4)). Over the past few decades, this expectation was amply confirmed by the successful mapping of neuronal activity in mesoscopic scale neuronal organizations using 7 tesla (T), initially orientation domains together with ocular dominance columns in the human visual cortex (5), followed by numerous and ever increasing number of submillimeter resolution human neuroscience applications (e.g., reviews (1,3,4,6)). Even at more conventional, supramillimeter resolutions typically employed at 3T and lower magnetic fields, 7T was shown to increase the detection sensitivity and model predictive power of fMRI (e.g., (7–9)), establishing the importance of ultrahigh magnetic fields (UHF, defined as ≥ 7T) for the study of human brain function irrespective of the spatial resolution and specific neuroscientific problems targeted. In addition, burgeoning activity in clinical applications has also demonstrated the presence of unique diagnostic advantages available at UHFs, particularly in the brain (review (10) and references therein). These accomplishments, together with the development of a plethora of technologies to overcome the challenges posed by imaging the human body at such high magnetic fields (e.g., (1,2,11)) has led to the introduction of commercially available clinical 7T MRI systems and, in parallel, catalyzed an interest in pursuing even higher magnetic fields for human imaging. An example of the latter is the 10.5T human MRI initiative undertaken in our laboratory (12–17).

The aforementioned UHF advantages arise mainly, though not exclusively, from magnetic field dependent increases in ultimate intrinsic signal-to-noise ratio (uiSNR), predicted by computational and theoretical works (18–23) and corroborated with experimental data (17,24). Capturing this maximum achievable SNR requires the development of complex radiofrequency (RF) coils. Such developments have largely taken place for 7T and lower magnetic fields, leading to complex arrays with multichannel transmit and receive capabilities for imaging of the human head (e.g., (25–36)). This, however, is uncharted territory for the new frontier of ≳10T imaging. Array designs derived from the 7T experience were recently shown to be suboptimal at 10.5T (24) and likely also at even higher magnetic fields. Another difficulty at UHF, due to correspondingly shorter wavelengths associated with the proton Larmor frequency, is that cables associated with the multitude of receive channels can couple to the circumscribing transmitter and act as scatterers (37). An interesting consequence of such coupling is the difficulty it creates in generating accurate electromagnetic (EM) models of the transmitter, a necessary step in defining safe operational limits.

In this paper, we describe the development of a 16-channel transmit and receive (Tx/Rx) array and a 64-channel receive (Rx) only array, which, when paired, create a 16Tx/80Rx configuration for brain imaging at 10.5T. This development incorporates engineering and computational solutions to the problem of determining safe operating limits and increasing the array performance towards capturing the available uiSNR. We report large field dependent SNR gains that can be attained at 10.5T relative to 7T with this array and present preliminary structural and functional images of the human brain obtained at 10.5T.

## 2. METHODS

### 2.1 RF Coils

#### 2.1.1. 10.5T Arrays

The 16-channel transmit, 80-channel receive coil assembly consist of two subassemblies: a 16-channel coil array with each element capable of transmit and receive functions (16Tx/Rx, i.e. 16-channel “transceiver” array) and a 64Rx only array.

The 16Tx/Rx array used self-decoupled (SD) elements (38) which are particularly suitable for the short wavelength at 10.5T. This design simplified the transmitter electronics and improved EM modeling (discussed further on); in addition, it exploits the hybrid dipole-loop (39) behavior of the SD concept and the favorable Poynting vector of dipoles (40–43).

The transmitter housing (Figure 1A), designed to be compatible with head gradient, was an open-ended cylinder, 37 cm long with inner and outer diameters of 27.5 and 32 cm, respectively. The SD concept enabled split-housings with no electrical connections between the two halves to improve workflow. Further details are provided in Supporting Information, Section 1.

**Figure 1:**
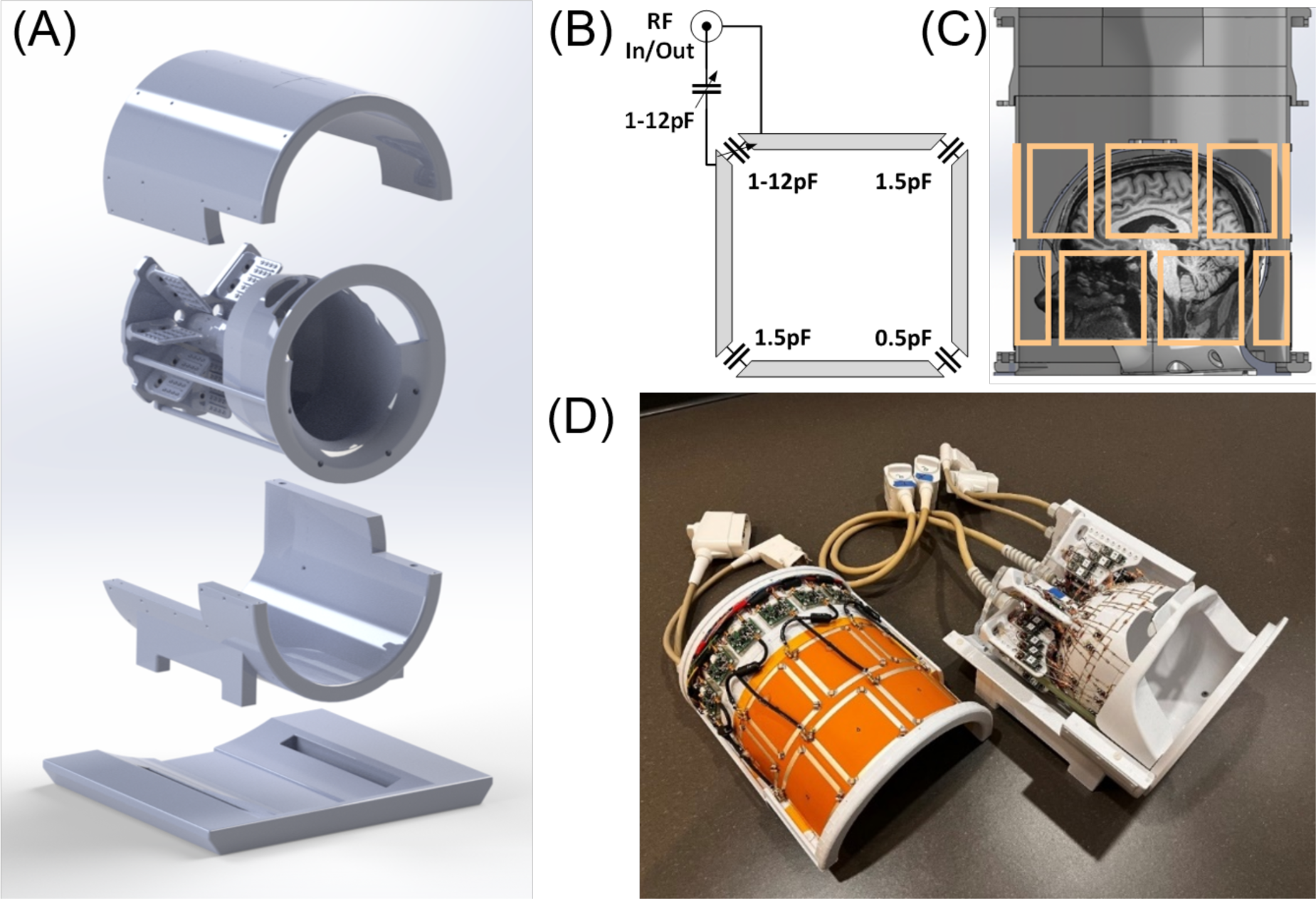
(A) 3D rendering of the 16-channel transmit, 80-channel receive assembly. (B) Schematic of a of single non-overlapping Self Decoupled (SD) transceiver element. (C) Representation of human head location within the receive-only insert positioned in SD transceiver array. (D) The complete 10.5T device assembly with the 16-channel SD T/R array and 64-channel receive-only array insert.

The 16 SD elements were tuned and matched at 447 MHz (10.5T ^1^H frequency) using a 16-channel vector network analyzer while loaded with a lightbulb-shaped phantom approximating the size and shape of the head and neck. This phantom was filled with a polyvinylpyrrolidone (PVP) solution with measured conductivity and relative permittivity of 0.65 S/m and 47.2, respectively, at 447 MHz (24).

Two versions of 16Tx/Rx arrays were fabricated (Figure 2). Both had the same element layout and circuitry; however, one (SD_e_) utilized an external MR system interface box containing transmit-receive (T/R) switches and Low Noise Amplifiers (LNAs) (SPF5122Z, Qorvo, Greensboro, NC, USA) connected to the coil elements with coaxial cables (RG400, Huber+Suhner, ^H^erisau, Switzerland) (Figure 2A). For the second (SD_i_), we designed and integrated a new miniaturized MR system interface into the array housing (Figure 2B), with integrated T/R switches (Figure 2C) on a 34×45 mm footprint, allowing them to be placed adjacent to one another azimuthally in the housing. Detailed interface notes can be found in Supporting Information, Section 2. The cumulative losses and phase lengths of the complete MR system interface were used to set up the initial B_1_^+^ shim for parallel transmission (pTx) operation, and ensure accurate modeling of our device in EM simulations.

**Figure 2:**
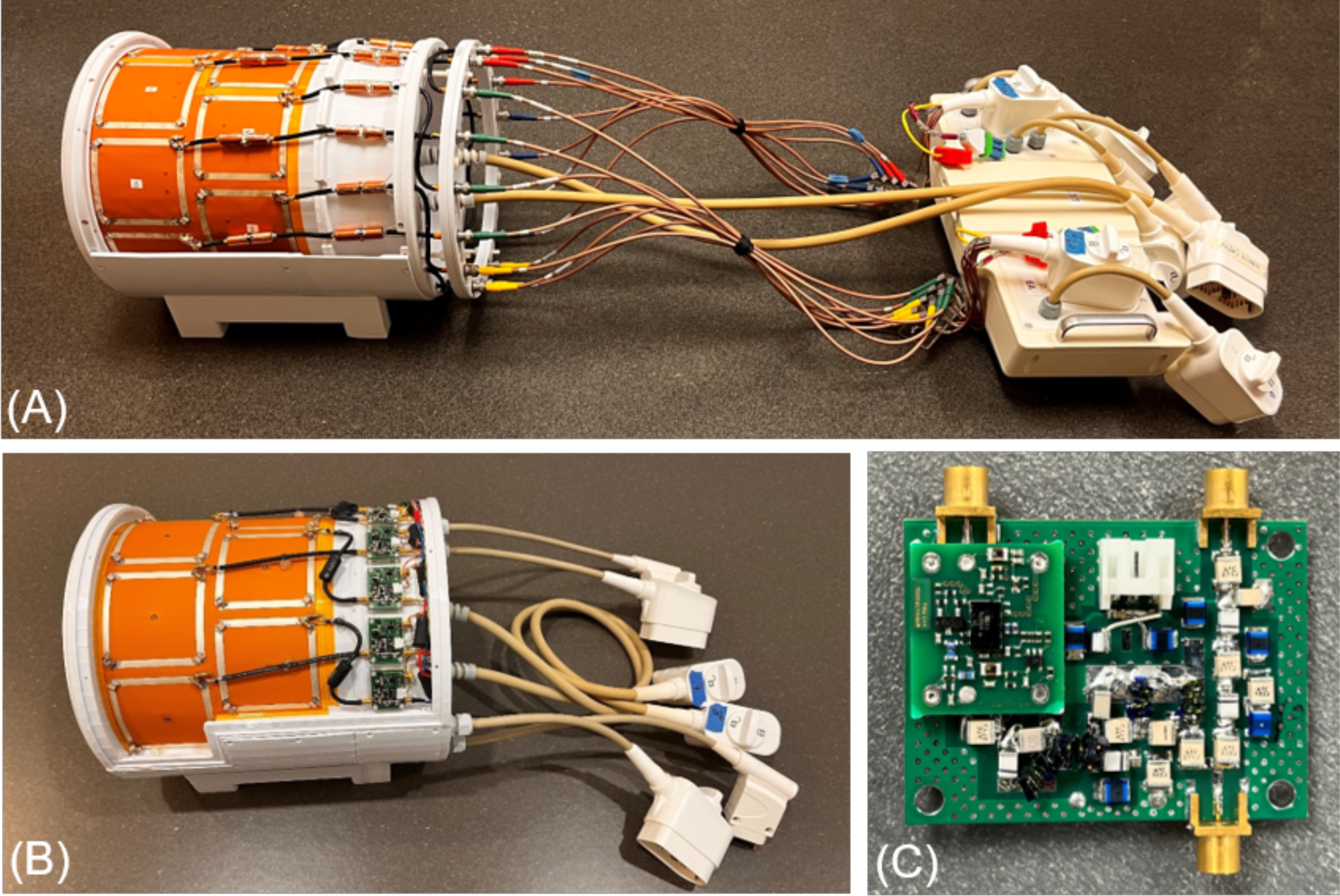
16 channel transceiver (i.e. SD_e_) configured to be used for transmit and receive operations using external T/R switches and preamplifiers (A), and a subsequent version (SD_e_)where a new miniaturized MR system interface containing T/R switches and a preamplifier was integrated into the array housing (B). The board containing the integrated T/R switches and the preamplifier are shown in (C).

The 64Rx array was designed to fit on a human head-conformal former which included open features over the nose and mouth. The inner surface of this former measured 22.5 cm (anterior– posterior) by 18.25 cm (left–right) with a wall thickness of 2.5 mm. The 64Rx array was laid out on the outer surface of the former and consisted of 64 loops in six rows along the z-axis (Figure 3D). The coil elements were overlapped in the superior-inferior direction, and non-overlapped azimuthally within each row. The majority of the elements were rectangular-shaped centrally (∼25×50 mm^2^) with both larger and smaller trapezoidal elements used to fit the tapered shape of the former in other places. Two larger elements (∼65×50 mm^2^) were employed over the eyes. Further details on the construction can be found in Supporting Information, Section 3.

**Figure 3:**
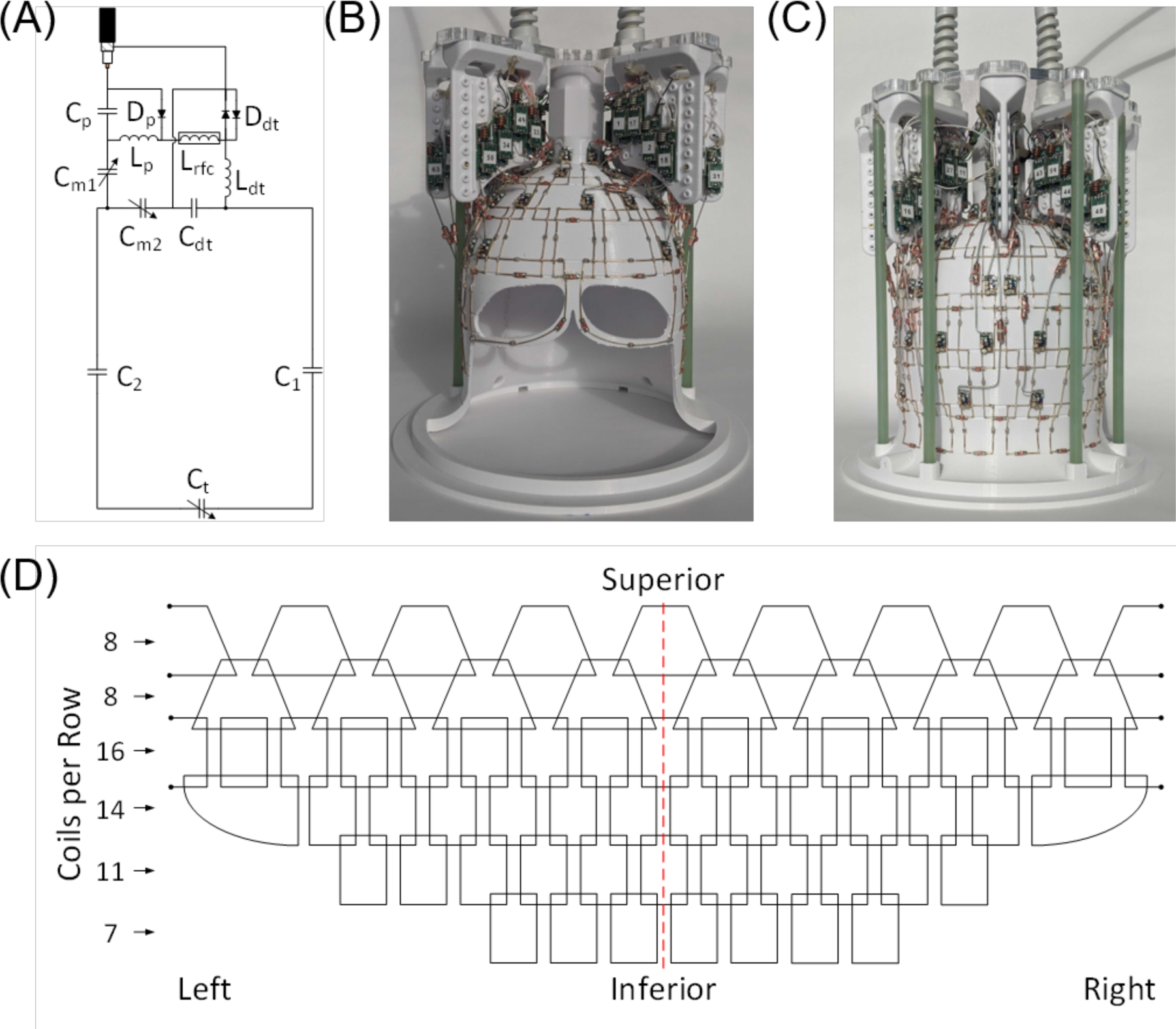
(A) 64-channel receive array element schematic. (B and C) the 10.5T 64-channel receive array as built. (D) Unwrapped 64-channel receive array layout (as viewed from the posterior). The red dashed line represents the posterior, while left and right ends would connect at the anterior. The number of coil elements per row are indicated on the left.

#### 2.1.2 7T 64Rx Receive-Only Array

A similar 64Rx array was fabricated for 297 MHz (7T ^1^H frequency). The geometry and layout of the coil elements matched exactly that of the 10.5T 64Rx array, though some component values were changed to adjust the resonant frequency. A 16Tx array with loop-like coil elements was used as a transmitter (44).

### 2.2 Electromagnetic Simulation and Safety Validation

The CMRR’s 10.5T scanner operates under an Investigational Device exemption (IDE) from the FDA and each RF coil must be FDA-approved prior to use with humans. A critical step in this process is the validation of the EM models developed for the arrays; this is done using phantoms, and in our case, the aforementioned lightbulb-shaped phantom mimicking the shape and average electrical properties of the human head and neck.

The RF-related patient safety evaluation of the 16Tx/80Rx array employed a 3-phase workflow (13,45). Phase one involved developing an EM model of the coil with good agreement between the experimental and simulated B_1_^+^-maps and S-parameters in the phantom. In Phase two, mismatch between experimental and simulated results were quantified and incorporated into the calculation of a safety factor (SF). Finally, this safety factor was used to scale the 10g-averaged spatially specific absorption rate (Q-matrices) derived from the EM simulations performed with human body models. The Q-matrices were eventually utilized to calculate the peak 10g-averaged spatially specific absorption rate (psSAR_10g_) for different RF excitation scenarios for determining safe power limits to comply with the IEC guidelines (46). Detailed safety validation notes can be found in Supporting Information, Section 4.

### 2.3 Data Acquisition

All volunteers for this study provided written informed consent. The study was conducted under an IDE from the FDA and approved by the University of Minnesota IRB, as described previously (14,15).

#### 2.3.1 RF Coils and MR Systems

All 10.5T data were acquired on a MAGNETOM (Siemens Healthineers, Erlangen, Germany) console interfaced to an 88 cm bore 10.5T magnet (Agilent Technologies, Oxford, UK) fitted with a Siemens SC72D gradient coil providing 70 mT/m maximum amplitude and 200 T/m/s slew rate. RF excitation employed 16 independent 2 kW RF power amplifiers (Stolberg HF-Technik AG, Stolberg, Germany) and signal reception utilized 80 of the 128 receive channels developed in house from Siemens components. The two 10.5T 16Tx/Rx arrays (SD_i_, SD_e_) used the same 64Rx insert. Individual complex B_1_^+^-maps for each of the 16Tx/Rx elements were acquired using the fast relative B_1_^+^-mapping technique (47) scaled utilizing the absolute B_1_^+^-maps acquired by actual flip-angle imaging (AFI) (48). These field profiles were used to generate pTx excitation patterns.

All 7T data acquisition was performed using a MAGNETOM Terra (Siemens Healthineers).

#### 2.3.2 Data for SNR

Images for SNR comparisons were obtained with a gradient-recalled echo (GRE) sequence with human (phantom) parameters of TR=10,000 (2,000) ms, TE=3.8 (3.5) ms, flip angle=90°, voxel size=2.0×1.0×2.0 mm^3^, 80 interleaved axial slices covering the whole brain (phantom). Identical noise images were acquired without an RF excitation pulse and TR=1,000 ms.

SNR was calculated following (25). uiSNR was calculated for the lightbulb phantom as described in (24).

#### 2.3.3. g-Factor and Parallel Imaging

The g-factor maps were calculated from the SENSE equations (49) for simultaneous multi-slice (SMS)/multiband (MB) acceleration with k-space undersampling (50–52), and for 1D and 2D k-space undersampling (25). The data for the g-factor maps were obtained with a fully sampled 1-mm isotropic 3D GRE sequence (25). For visualization, maximum intensity projections (MIP) of g-factor maps were extracted in the left-right (LR) direction over an 80-mm thick slab in the sagittal plane and displayed over a silhouette of the central sagittal slice as 1/g maps. The SNR of the accelerated images (*SNR*_*acc*_) was calculated as:

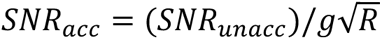

where *R* is the reduction factor for k-space undersampling, and *SNR*_*unacc*_ is the SNR without acceleration (49).

#### 2.3.4 Anatomical Imaging

Anatomical images were obtained with GRE, MP2RAGE, and SWI sequences. Acquisition parameters are given in the figure captions.

GRE and MP2RAGE anatomical data were obtained in the 16 channel pTx mode using phase and amplitude shimming; SWI images used a circularly polarized mode.

#### 2.3.5 Functional Imaging

fMRI with visual stimulation was acquired using 0.8 mm isotropic 3D Echo Planar Images (EPI), with a single shot to cover the k_x_-k_y_ plane (iPAT = 5) and segmenting along k_z_ using TR=68 ms, TE=18.2 ms, Volume Acquisition times (VAT)=3,808 ms, Partial Fourier=6/8, and flip angle ∼15°. Detailed processing and stimuli are described in Supplementary Material, Section 5.

## 3. RESULTS

### 3.1 Bench Measurements

#### 3.1.1 16-channel Tx/Rx Array

Bench measurements demonstrated good tuning and matching for the 16Tx/Rx array elements (SD_i_, SD_e_), for both the phantom load and the human head, with and without the 64Rx insert. The return losses (coil matching values, S_ii_, for the phantom-loaded 16Rx/Tx array with the integrated T/R switches and preamplifiers (SD_i_) and the 64Rx array in place ranged from -11.8 dB to -35.6 dB, with an average of -20.9 dB, while the inter-element coupling (S_ij_) ranged from -9.4 dB to - 48.7 dB with an average of -27.5 dB; SD_e_ demonstrated comparable values. Most of the coupling within the 16Tx/Rx arrays happened between the diagonal neighbors, which ranged from -9.4 dB to -22.7 dB, with an average of -15.3 dB, and was sufficient to permit tuning of each individual element with minimal interaction. Q-ratios, evaluated as Q_HCSS_/Q_L_ were ∼3:1 when measured with a decoupled probe pair (53).

#### 3.1.2 Integrated MR System Interface

In the transmit mode, an average insertion loss of -0.4 dB with 0.7° phase balance was measured for the 16 integrated T/R switches and preamplifiers of SD_i_ (Figure 2B); the preamplifiers were isolated from the transmitter-to-coil path by 50.1 dB, protecting them during excitation. The two 8-channel pTx plug assemblies introduced an additional 0.55 dB loss in the transmit path, accumulating to ∼1 dB of total insertion loss from the system connection to the coil element. The preamplifiers had a gain of ∼28.0 dB, noise figure (NF) of 0.45 dB, and insertion losses in the receive path preceding the preamplifier measured -0.59 dB; the transmitter input port was isolated from the receive path by 49.3 dB.

The external interface box used in conjunction with SD_e_ (Figure 2A) demonstrated an average of -1.1 dB insertion loss and phase balance of 6.6° across the 16 channels; ∼0.54 dB of additional loss in the transmit paths was introduced by the coaxial cables and system adaptors. The LNA integrated into the external interface box produced ∼20.0 dB of gain with a NF of ∼0.5 dB at 447 MHz; the TR switches added insertion losses of ∼0.5dB into the receive path prior to the LNA.

#### 3.1.3 The 64-channel Receive-Only Array

All elements in the 10.5T 64Rx array were matched to -15 dB or better with the phantom described above. It was impractical to measure the complete S-parameter matrix of this array; however, simulations for the 25×50 mm^2^ non-overlapped neighboring elements (azimuthal direction) demonstrated −11.2 dB coupling (S_21_) compared to −13.8 dB for the overlapped elements (superior-inferior direction); for the 50×50 mm^2^ elements, the non-overlapped neighbors exhibited −12.9 dB coupling compared to −16.8 dB with the overlap. This degree of isolation was considered sufficient given that preamplifier decoupling (54) further improves the isolation. Among the multiple-sized array elements, the Q_U_ to Q_L_ ratios ranged from 7.9:1 for heavier loading conditions to 2.2:1 for lighter loading conditions. The 64Rx elements demonstrated typical active PIN detuning of ∼37 dB during excitation. Similar data are provided for the 7T 64Rx array in Supplementary Information.

### 3.2 Simulation Results and Safety Validation

The simulated (without the 64Rx) and experimental (with the 64Rx) S-parameters for the 16Tx/Rx transceiver are presented in Supplementary Figure S2, displaying good agreement. The experimental and simulated B_1_^+^-maps corresponding to the CP mode of excitation are given in Supplementary Figure S3. The agreement between the simulated and experimental B_1_^+^-maps were inferior to the excellent agreement we previously achieved for a 10.5T 8-channel transceiver array (13). This is expected since the 64Rx insert is not altogether transparent to the transmitter (Figure 4) despite efforts to minimize the interactions; for this excitation pattern (CP mode), a 25% normalized root-mean-square error (NRMSE) between the simulated and measured B_1_^+^-maps was calculated inside the uniform, lightbulb-shaped phantom for the 16Tx/Rx array with the 64Rx insert in place. This discrepancy required the calculation of the EM modeling error, *e_EMM_* ; the error between simulated and measured B_1_^+^-maps was propagated to the peak-SAR_10g_ error using 10^6^Monte-Carlo simulations. Supplementary Figure S4 shows the histogram of the propagated SAR error for the CP mode. The *e_EMM_* for this coil was calculated as 48%, which yielded an SF of 1.71, assuming an inter-subject variability, *e_ISV_* (55), of 50% and a power monitoring uncertainty, *e_PM_*, of 15%. Using the VOPs scaled up by the safety factor, a total input power limit of 22 W was calculated for the CP-like excitation in the first-level operating mode in compliance with IEC guidelines (46).

**Figure 4:**
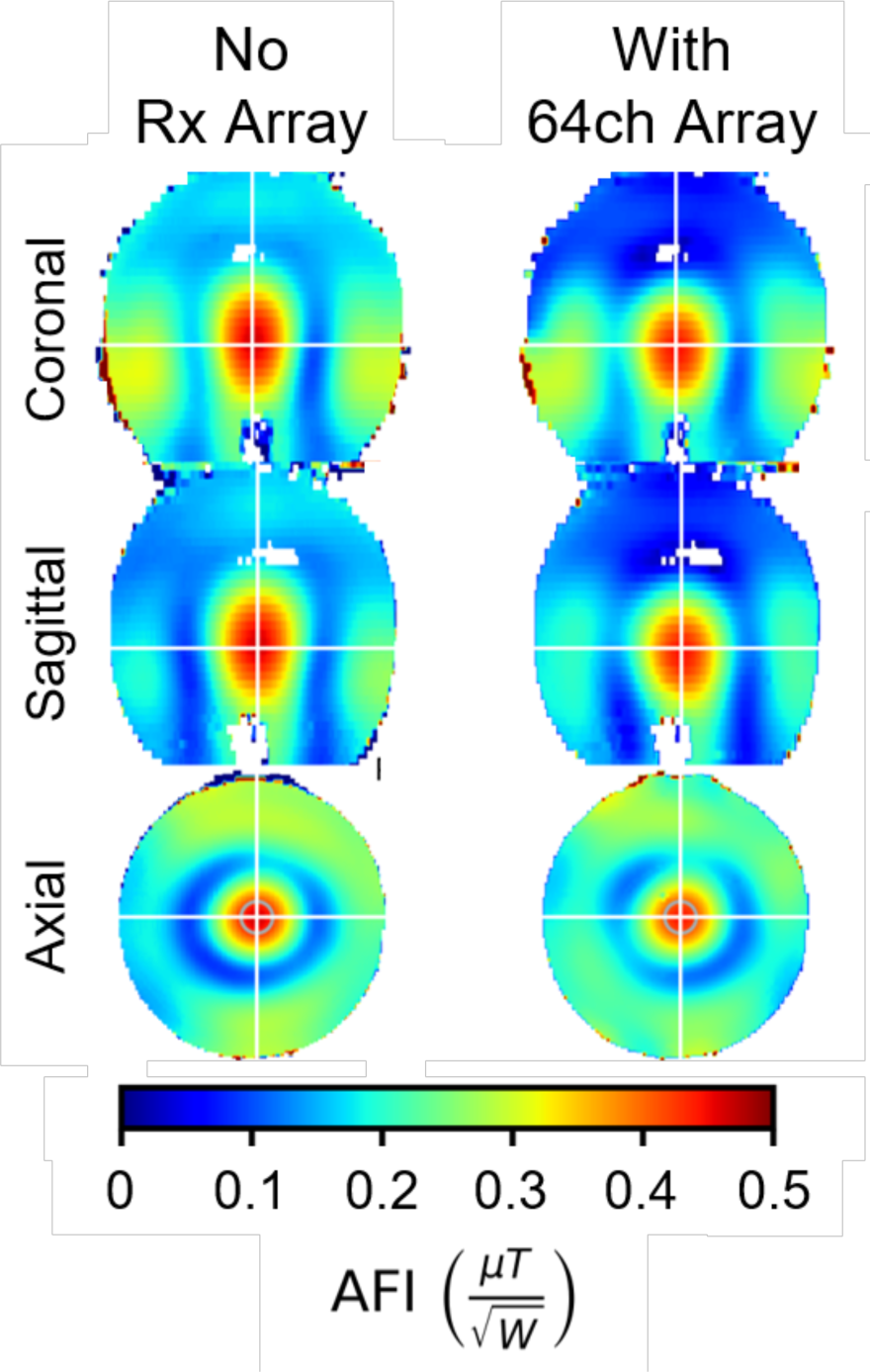
Experimentally measured AFI maps of the 16-channel SD array (A) without the 64-channel receive-only insert, and (B) with the 64-channel receive-only insert.

### 3.3 Experimental Results

#### 3.3.1 SNR versus Array Configuration

Combining the receive capability of the 10.5T 16Tx/Rx array (SD_i_) with the 64Rx insert created an 16Tx/80Rx configuration. Supplementary Figure S5 shows the 80Rx noise correlation matrix, demonstrating an average value of 5% with a phantom load and 4.5% with a human subject.

Figure 4 illustrates AFIs from the 10.5T 16Tx/Rx SD_i_ array with and without the 64Rx insert in place. A mean of 0.440 and 0.419 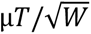 was achieved inside of a centrally located 314 mm^3^ region of interest (ROI) (denoted by the blue circle in Figure 4, axial slice) with and without the 64Rx insert.

SNR gains at 10.5T were observed throughout the phantom as a result of integrating the system interface into the coil housing (Supplementary Figure S6). SNR was quantified by averaging it in 1 cm thick concentric shells, with the outer-most shell conforming to the boundaries of the phantom (Figure S6B); using this approach, a one-dimensional plot showing SNR against depth of shell, was generated (Figure S6C), demonstrating SNR gains of ∼5-10% in the periphery and ∼25% near the center of the phantom where the SNR is lower.

The 80Rx configuration yielded SNR gains relative to the 64Rx insert alone throughout the human head (Figure 5) and the phantom (Figure 6) at 10.5T. These gains are most pronounced in the central regions where the SNR is intrinsically low and in regions of reduced coverage by the 64Rx array. Figure 6 shows the SNR gain also as a one-dimensional plot obtained using the afore-described 1 cm thick concentric shells. As much as ∼40% gain is observed centrally, and ∼5 to 10% peripherally, where the SNR is intrinsically high. Additional demonstration of these gains are provided in Supplementary Figure S7.

**Figure 5:**
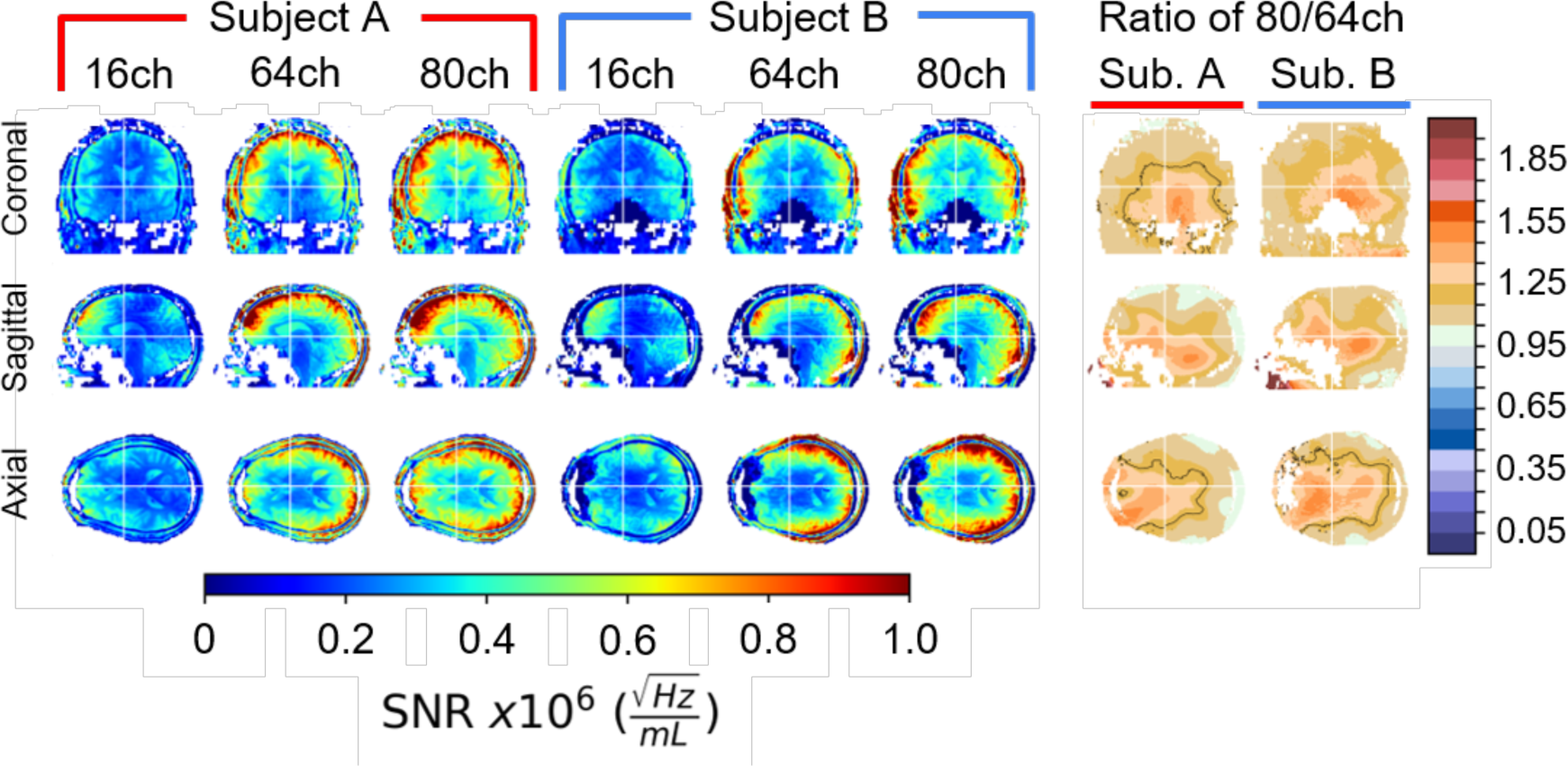
In-vivo SNR gains achieved at 10.5T using the 64ch receive-only array *versus* the combined 80ch receive assembly. SNR data from the 16 SD array, 64ch receive-only array, and the combined 80ch receive assembly are shown for two different human subjects in the left 6 columns. The right two columns show ratio maps of the combined 80ch receive vs the 64ch receive array, demonstrating nice global improvements in SNR with a central boost of ∼50% from the additional 16 SD channels.

**Figure 6:**
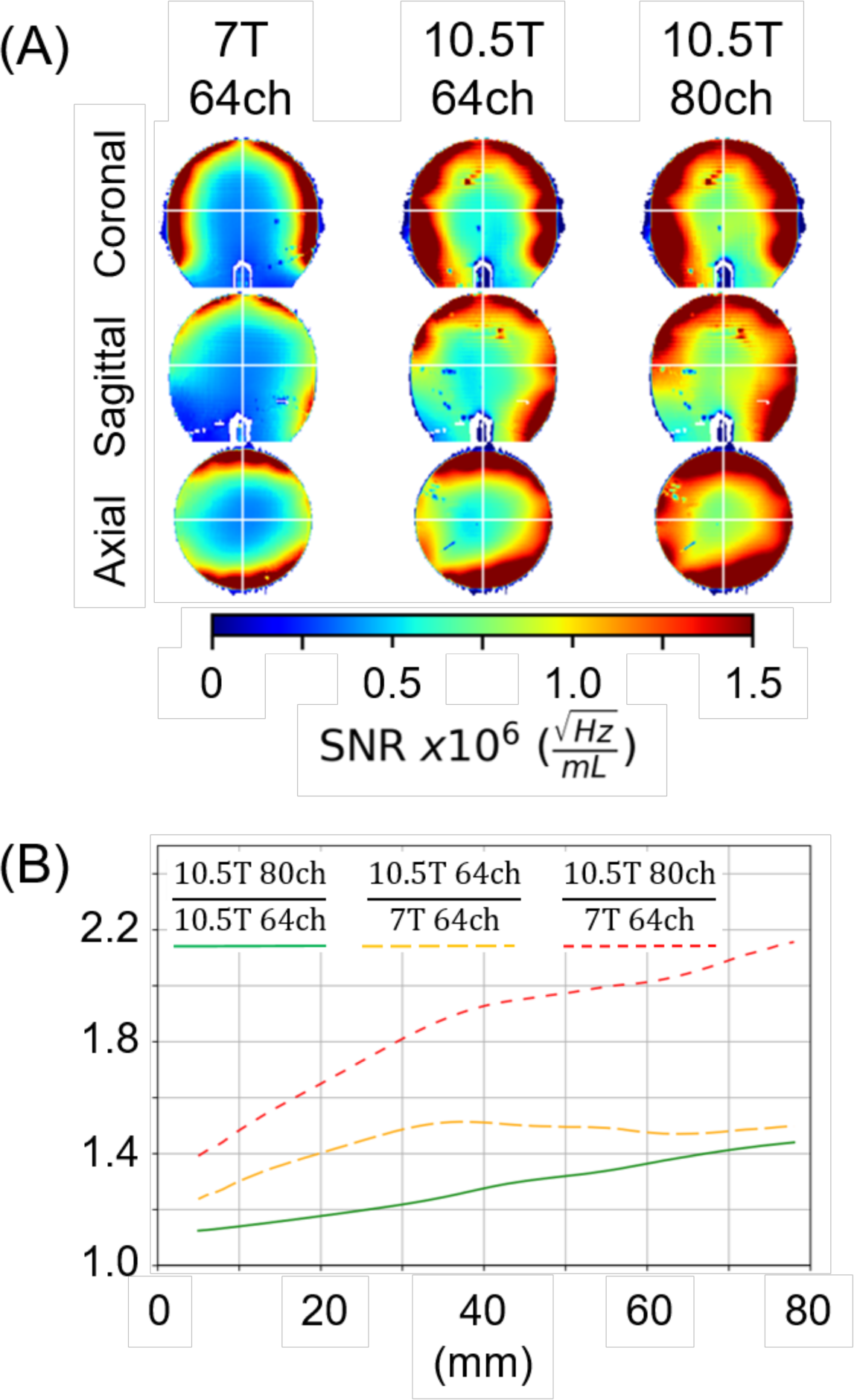
SNR gains related to magnetic field strength and array configuration measured with the “light-bulb” shaped phantom. Panel (A) displays SNR maps for the 64ch receive-only array at both at 7T and 10.5T (left-most and middle columns, respectively), and for the 80 channel receive configuration at 10.5T (right most column) for three orthogonal central slices. (B) The ratio of SNR as a function of depth plotted for SNR averaged in 1 cm thick concentric shells. Red-dashed curve is the ratio of SNR of 10.5T 80Rx to the SNR of 64Rx 7T. Yellow dashed curve is SNR ratio of the 64Rx array at 10.5T versus 7T. The green curve is the SNR ratio of the 80Rx versus 64Rx both at 10.5T.

#### 3.3.2 SNR Across Field Strengths, 7T vs. 10.5T

SNR gains measured in the lightbulb phantom for the different array configurations, corrected for different T_2_* at the two field strengths are shown Figure 6; SNR maps in three central and orthogonal slices are illustrated (Figure 6A) demonstrating clearly visible gains for the identical 64Rx arrays in going from 7T to 10.5T and to the 80Rx configuration at 10.5T. Figure 6B shows this SNR gain as a one-dimensional plot using the previously described 1 cm thick shells. Relative to the 7T 64Rx array, the identically laid out 10.5T 64Rx array by itself achieves ∼1.5-fold SNR gain. In contrast, the 10.5T 80Rx configuration demonstrates a factor of ∼2 SNR increases over a large central volume relative to the 64Rx array at 7T, reaching ∼2.2 in a small central region.

Figure 7 shows performance maps (56) that display the SNR in the lightbulb phantom at 7T and 10.5T as a percentage of the corresponding uiSNR. In agreement with our previous report (24), 63Rx or 64Rx arrays with identical layouts at 7T and 10.5T capture a lower fraction of the available uiSNR at 10.5T compared the 7T (∼53% vs. ∼76%). On the other hand, as expected from Figure 6, 80Rx increases the fraction of uiSNR captured at 10.5T to ∼71% but still remains lower than that achieved by the 64Rx array at 7T.

**Figure 7:**
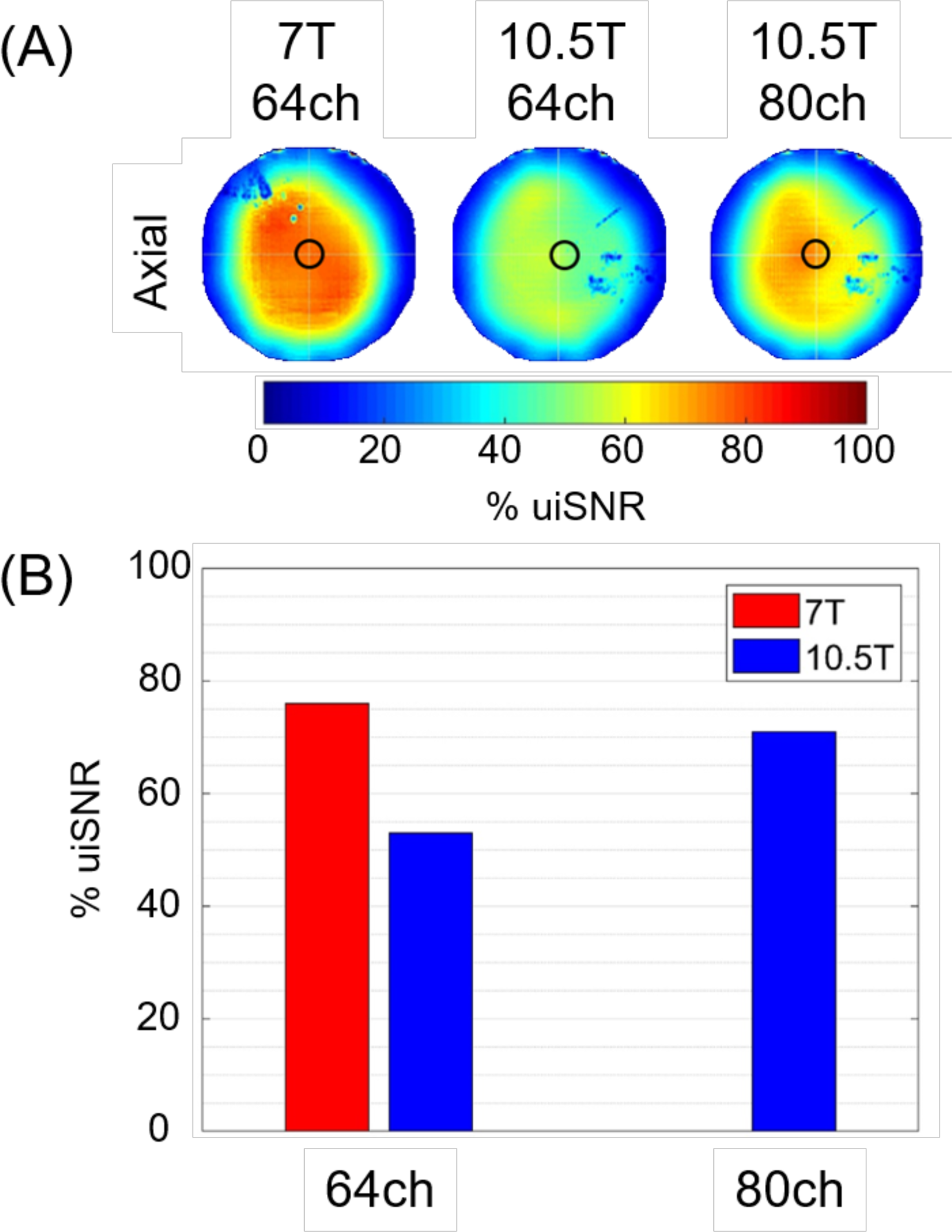
The SNR of the arrays as a ratio of the uiSNR, presented as a map in a central axial slice of the lightbulb shaped phantom (A), and as measured in small 2cm diameter central spherical volume, identified by a black circle in panel A, presented as bar graphs (B). uiSNR is different for 7T and 10.5T and was calculated separately for the two field strengths.

#### 3.3.3 g-Factor and Parallel Imaging

Since realistic image acquisitions employ parallel imaging, we evaluated the g-factor for the two 10.5T coil configurations for k-space undersampling (49) and/or the use of slice acceleration by the SMS/MB technique (50–52,57,58).

Figure 8A displays 1/g as a maximum intensity projection (MIP) onto a sagittal slice for a central 80 mm slab for 3 different acceleration patterns, demonstrating that the additional 16 channels from the transceiver actually marginally degrades the g-factor noise. The effect of this g-factor noise on the SNR for the same three accelerated acquisition patterns for both coil configurations is shown in Figure 8B, which plots 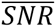 (i.e. SNR_unacc_/g) for a single central 1mm think sagittal slice. Figure 8B demonstrates that the gains in unaccelerated SNR makes up for more than the slight degradation in the g-factor noise so that 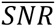 is higher for the 80Rx configuration. This last point is further illustrated by presenting maps of 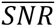 as a ratio for the 80Rx *versus* the 64Rx configurations in Figure 8D over the central 80mm slab.

**Figure 8:**
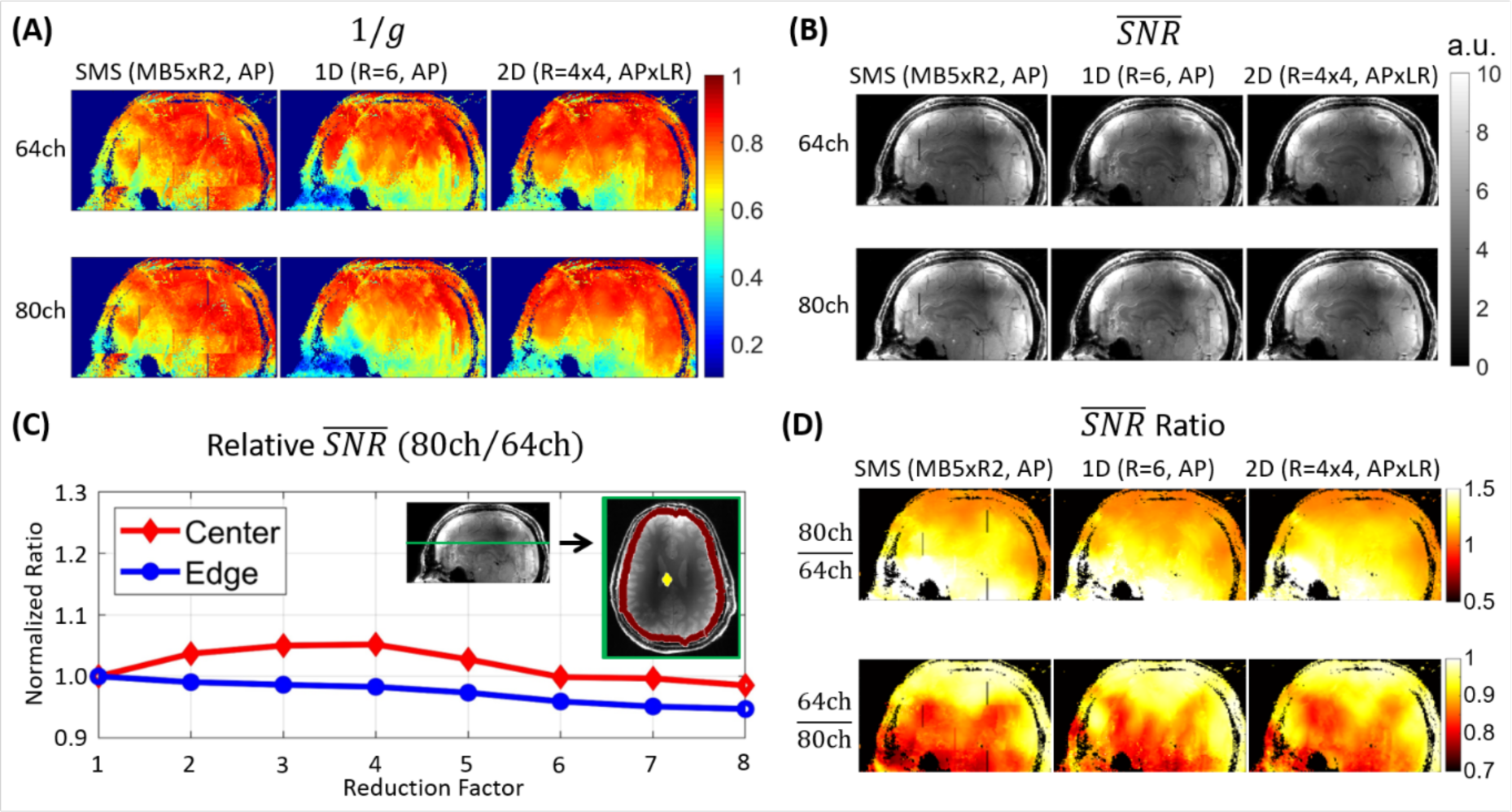
(A) 1/g map for different undersampling strategies and channel configurations. The leftmost column shows 1/g for SMS/MB acquisition for MB=5 and in-plane reduction factor R of 2 (i.e., MBxR=5×2) in the superior-inferior (SI) and anterior-posterior (AP) directions, respectively. The middle and right most columns in this panel show 1/g for 1D acceleration with R=6 (AP) and 2D 4×4 (APxLR) accelerations, respectively. The top row shows the 1/g MIP maps with the 64-channel receiver array where the MIP is over an 80 mm central slab. The bottom row shows the corresponding maps for the 80 channel configuration. (B) 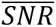, defined as (*SNR*/*g*), maps for different undersampling strategies and channel configurations. The top row shows the 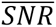maps with the 64-channel receiver array for SMS, 1D, and 2D undersampling, and the bottom row shows the corresponding maps for the 80-channel configuration. (C) 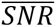 improvement for the 80-channel configuration relative to 64-channel configuration for different phase-encoding undersampling values. The anatomical image is segmented into an edge ROI and central ROI, and in the line-plot the average difference in SNR between the coil configurations is plotted for each undersampling factor. (D) The MIP of the ratio in 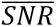 between the 80- and 64-channel coil configurations for the 3 different acceleration scenarios.

In addition, we examined the impact of different k-space undersampling factors (*R*) factors by considering 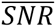 of the 80Rx configuration relative to the 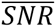 of the 64Rx array (Figure 8C) for the 1D acceleration scenario as a function of *R* for a central ROIs and a peripheral ROI defined as an ∼1 cm band circumscribing the edge of the brain. The mean 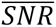 ratio for the different ROI’s, plotted for different *R* factors, remained stable over the large R factor range.

### 3.4 In vivo Imaging

#### 3.4.1 Anatomical Images

Figure 9 shows two axial (top row) and sagittal (middle row) slices from 0.2×0.2×1 mm^3^ GRE images, acquired in ∼5 min, illustrating clear visualization of several different brain structures: white and gray matter, different nuclei within the thalamus, optic radiation emanating from the lateral geniculate nucleus and projecting into the visual cortex, venous blood vessels including intracortical radial ones, and the line of Gennari, a group of myelinated axons that run horizontally along the middle of the cortex in primary visual area V1 (particularly prominent in the sagittal slice shown on the left panel); this canonical feature of V1 is visible on both sides of the calcarine fissure, as expected, since V1 is located in both the lower and upper banks of the calcarine fissure. Additional slices, and expanded and annotated versions of these images are provided in Supplemental Figure S9.

**Figure 9:**
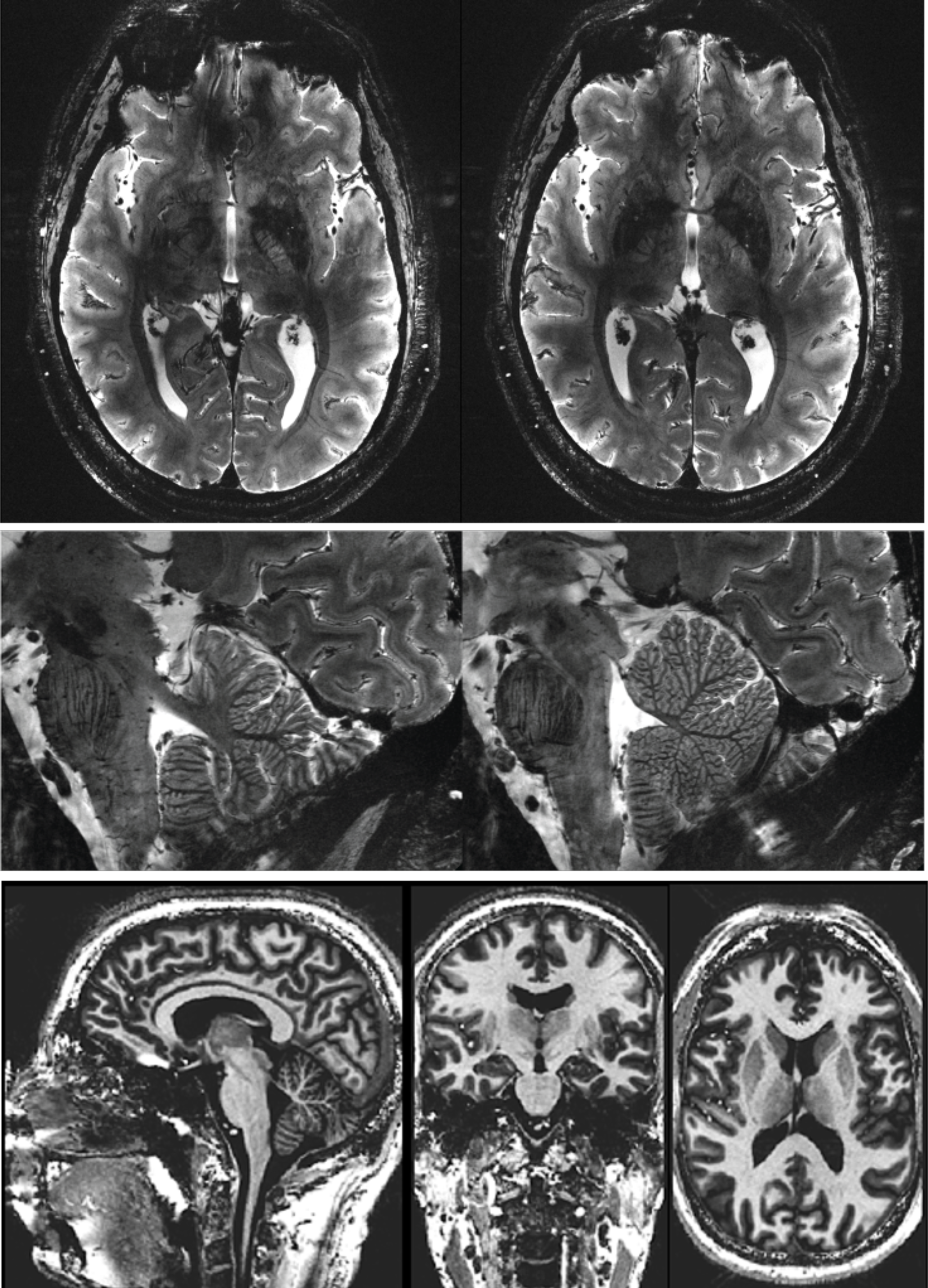
Example of anatomical images at 10.5T obtained with the 16Tx/80Rx configuration. Top and middle rows: T2* weighted GRE Images acquired with 0.2×0.2mm in plane resolution with 1.0 mm slices in the axial (Top row) and sagittal (middle row) orientations, displaying exquisite detail including the line of Gennari, particularly visible in the axial and sagittal images on the left. The images were acquired in 4.49 and 5.28 min in the axial and sagittal orientations respectively (TR=600ms; TE=20 ms; Nominal FA = 35°; in-plane acceleration factor (iPAT) = 2 (for the axial images only); bandwidth = 40Hz/pixel; Matrix size = 1024×896 for the axial and 1024×1024 for the sagittal images) Bottom Row: MP2RAGE Images demonstrating whole brain coverage, MP2RAGE Imaging parameters were: 0.7 mm isotropic voxels; TR = 5,000 ms; TA = 7.5 min; TE = 1.970 ms; FoV 253*270 mm^2^; in plane acceleration factor (iPAT) = 2; Inversion 1 = 840 ms; Inversion 2 = 2370 ms; partial Fourier = 6/8. All images were acquired in the pTx mode using phase and amplitude B1 shim.

The bottom row of Figure 9 shows MP2RAGE images acquired in 7.5 min, illustrating the ability to achieve whole brain coverage, tolerance to 180° pulses, and the ability to achieve spatially uniform transmit B_1_ necessary to impart spatially uniform contrast all within operational SAR limits at 10.5T.

#### 3.4.2 Functional Imaging

Figure 10 shows 10.5T fMRI data for visual stimulation with a face perception task in a human subject obtained using 3D GRE EPI, with nominal 0.8 mm isotropic resolution. These are the first human fMRI results at such a high magnetic field. Representative slices from a single 3D EPI volume are shown. These slices cover regions of the brain that are typically difficult to image with EPI due to B_0_ inhomogeneities induced by air filled cavities (Figure 10A). Figure 10B displays functional maps in the visual cortex in 3 different orthogonal planes, presented without smoothing and generated *without* the use of a denoising technique, such as NORDIC (59). Single voxel time courses are shown in Figure 10C.

**Figure 10:**
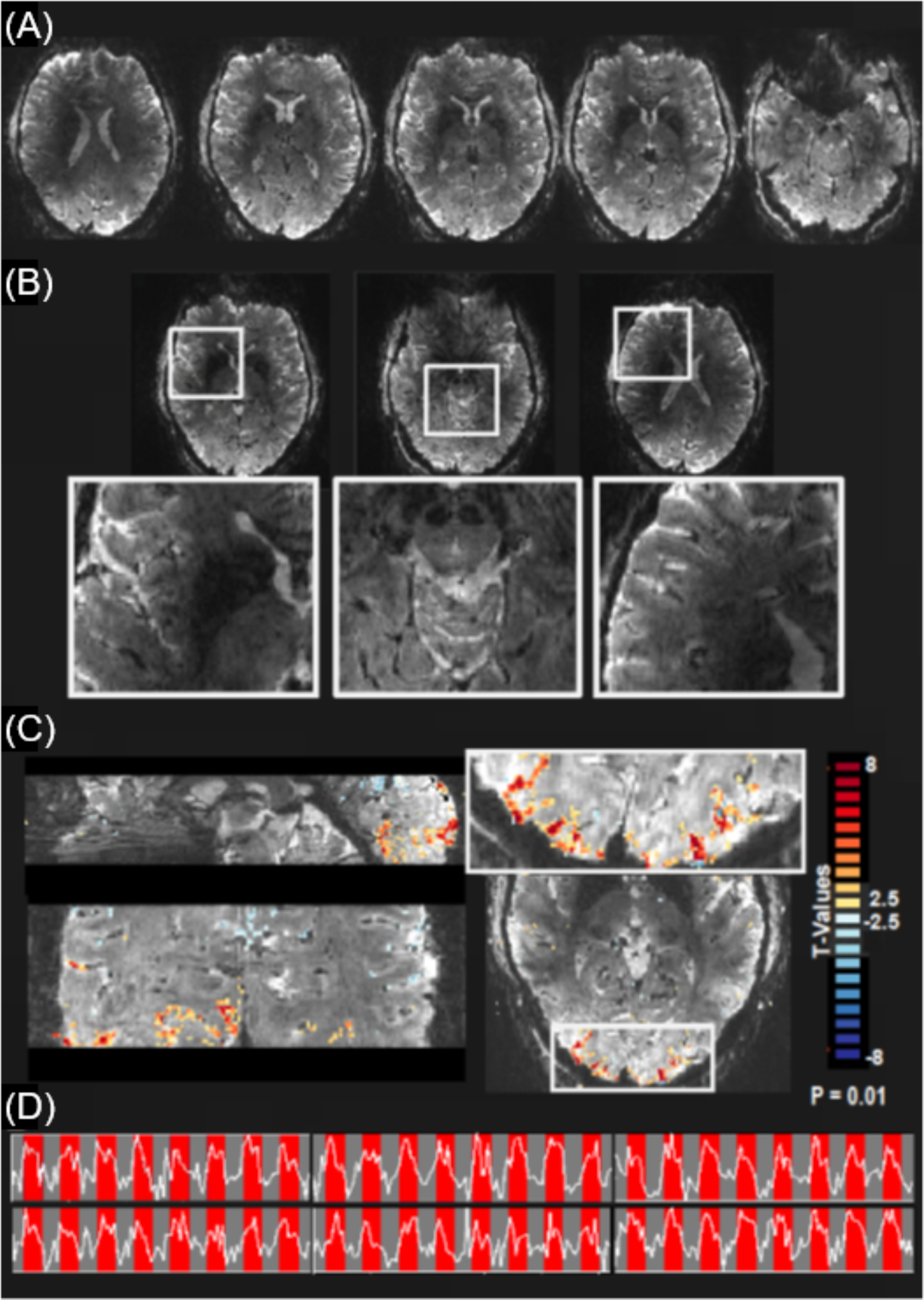
(A) An example of individual slices from a single EPI volume covering the whole brain. Selected slices in panel A illustrate the large coverage in this EPI acquisition including regions near the nasal cavities which are highly problematic for EPI acquisitions. (B) Observed t-maps displayed in 3 orthogonal orientations computed for the contrast faces > 0, superimposed on single EPI slices, with a zoom in view (green box) of the occipital cortex. (C) Single voxels time courses; red bars indicate stimulus presentation window, while the gray bars indicate baseline periods.

## 4. Discussion

### Transmitter Design

We successfully designed a 16-channel transceiver array relying only on self-decoupling principles and evaluated the combined use of this transceiver with a 64-channel receive-only array at 10.5T.

The self-decoupled design was primarily motivated by the need to increase the accuracy in the EM model for determining the safe operational power limits. In our previous work with a 10.5T 8Tx/Rx head array (13), the strategy employed was to replace all lumped elements by excitation ports, similar to the co-simulation method (60), and subsequently minimize the RMS error between simulated and measured complex S-matrices. This strategy produced EM model predicted B_1_^+^ and SAR maps that were in excellent agreement with experimental data under a variety of conditions. However, with the first 16Tx head array we developed to be used together with receive only inserts, this approach yielded suboptimal results leading to a large modeling error. The initial 16Tx array used conventional loops segmented by multiple capacitors and was inductively decoupled (Supplementary Figure S8); otherwise, the layout was similar to the current SD 16Tx/Rx array. One contributor to this large modeling error with this first transmitter, though not the only one, was the size of the “port-space” over which the optimization was performed: 24 for the 8Tx/Rx array *vs*. 400 for the inductively decoupled 16Tx array. Going to the SD loop design, which was easy to develop for 10.5T due to the shorter wavelength, decreased the “port-space” dimension to 64. The resulting EM model, although not perfect, was substantially improved and led to practical safety limits on the power utilized, allowing, for example, whole brain 90° and 180° flip angles in power demanding sequences like MP2RAGE (Figure 9). Importantly, this new transceiver coil supported a splitable housing with no electrical contacts (Figures 1C, 2B), making it easier to position a variety of receiver coil arrays, subjects, as well as fMRI and positioning apparatus.

The identical 64Rx arrays captured a much larger fraction of the central uiSNR at 7T compared to 10.5T (Figure 7 and also see (24)); this implies that there remains room for improvement. Simulations suggested that simply increasing the 64Rx loop sizes so that the layout utilizes fully overlapped loops did not alter the performance significantly (data not shown). An alternative approach was to employ the 16-channel SD transmitter as a transceiver, a configuration we had employed previously (34,35). The recent extension of our 10.5T frontend to a full 128 channel receiver system allowed us to explore this 16Tx/80Rx configuration with an integrated MR system interface to reduce the overall losses in the transmission path and improve SNR (Supplemental Figure S6). We attribute this gain to the cumulative benefits of shorter coaxial cables before the first amplification stage and improved inter-element decoupling during acquisition.

### Field dependence of the Signal-to-Noise Ratio

The data presented here and in our previous paper (24) highlight the difficulty of quantifying the field dependence of SNR at UHFs; at these very high magnetic fields, array coils composed of loop elements with identical layouts are not equally efficient at tapping into the available uiSNR (Figure 7 and (24)); a similar conclusion was reached in another numerical simulation study (61) where it was demonstrated that the central SNR performance of a 64-channel loop receiver drops to 80% at 11T while it is nearly 100% at 7T. Therefore, field dependence of SNR can be inferred only under conditions where the discrepancy with respect to the ability to capture the uiSNR does not exist or it is accounted for.

We have previously reported that, in going from 7T to 10.5T, uiSNR calculated in a human head model increases 1.4-fold peripherally, 2-fold at the edge of the brain, and a 2.6-fold centrally (24), with similar increases calculated for the lightbulb phantom. In this paper, we demonstrate that we come close to capturing this predicted SNR gain with the 16Tx/80Rx configuration. As measured in the lightbulb phantom, we observe ∼2-fold gain in a large central region with the 80Rx array at 10.5T compared to the 64Rx array at 7T, with the gain increasing to ∼2.2-fold in the very center (Figure 6). However, these numbers should be corrected by the fact that the 10.5T 80Rx and 7T 64Rx coils capture ∼71% and 76% of the uiSNR calculated for this phantom, respectively. This would translate into an SNR gain relative to 7T of ∼2.2 in a large central region and a ∼2.35 in a smaller central region. These gains are also consistent with our previous work conducted in collaboration with Neurospin and Maastricht colleagues, where SNR in the center of a sphere was reported to increase as 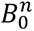 where *n* ≈ 2 (17); however, this conclusion was valid only in a small volume-of-interest (VOI) in the very center of the spherical phantom where B_1_^+^ and B_1_^-^ fields from all current carrying elements of the birdcage coils employed in that study add constructively (62); outside of this small central VOI, SNR in fact decreases with increasing magnetic fields in this setup (17), as expected due to destructive interferences (62). Unlike this previous work, we demonstrate here a practical array solution that approaches the uiSNR over a large central volume and, at the same time, provides additional advantages like parallel imaging.

Nevertheless, even this 80Rx configuration leaves room for improvements since it only captures 71% of the available uiSNR; further gains could be possible, for example, with a 128Rx array, which we have reported so far as an abstract (63). Different coil types, particularly dipole like antennas in addition to loops, could also play an important role as; this was predicted with previous analytic studies (43), and also demonstrated experimentally with 32 element arrays at 10.5T (64) and 7T (30,65).

Such field-dependent SNR gains are critical in exploiting the potential of ultrahigh magnetic fields in brain imaging. Particularly important are the gains centrally, where the uiSNR is inherently low. At times, it is argued that high numbers of small coils tiled over the head provide sufficient SNR gains to enable the study of the cerebral cortex. However, the cerebral cortex is not just a layer approximately equidistant from the brain-skull interface and the elements of head-conforming receiver arrays; rather, it is highly convoluted and reaches all the way into central regions of the brain (e.g., see Figure 9). Therefore, the field dependent SNR gains centrally are critically important for the cortex as well as subcortical nuclei (66).

In peripheral regions, such as the skull-brain interface, the uiSNR is inherently very high and thus, arrays capture a far smaller fraction (67). In this regime, increasing the number and density of elements in the array together with higher magnetic fields should provide improved performance (67).

### Anatomical and Functional Images

We demonstrated the utility of the 16Tx/80Rx array at 10.5T for anatomical and functional imaging (Figures 9 and 10, respectively). These images are not meant to explore the ultimate limits of resolution and/or contrast available at 10.5T; they are preliminary results illustrating feasibility and potential.

Higher magnetic fields provide increased T_2_* contrast; together with achievable high resolutions, this leads to impressive depictions of many structures (Figure 9, top and middle rows). We also demonstrate that whole brain T_1_ weighted images can be obtained at this very high magnetic field (Figure 9, bottom row). The T_1_ contrast is maintained at 10.5T as expected based on our earlier work (68). The use of parallel transmission enables uniform T_1_ contrast, which is otherwise difficult to achieve in the presence of transmit B_1_ inhomogeneities.

Finally, we demonstrated whole brain BOLD fMRI using GRE EPI at 0.8 mm isotropic resolution. Difficulties in obtaining good EPI images at high magnetic fields are well known. However, higher accelerations attainable with the synergistic use of high channel count arrays and high magnetic fields ameliorate such difficulties. Although distortions are visible due to high B_0_ inhomogeneities near the nasal cavities, these are not worse than for typical 3T EPI images. Robust functional responses and single voxel time courses where the stimulus induced changes can be visualized with ease are also presented (Figure 10).

The functional data presented here were not denoised using algorithms like NORDIC (59). Denoising will enable further significant improvements. In addition, particularly the EPI images will benefit from high performance gradients with high slew rates that are under development for 10.5T.

## 5. Conclusion

Dedicated multichannel transmitters paired with high channel count receive arrays are critical for exploiting UHF advantages. Here we explore, for the first time, the efficacy of such multichannel transmit and receive array configurations at the uniquely high magnetic field of 10.5T. We evaluate coil performance in the context of uiSNR and demonstrate that array designs composed of loop elements developed for 7T do not translate well to 10.5T. We also present solutions that remedy this problem and capture a large fraction of the theoretically maximal SNR available at 10.5T; anatomical and functional imaging data obtained with such SNR gains illustrate the feasibility and exciting potential of human brain imaging at magnetic fields greater than 10T.

## Acknowledgments

We dedicate this work to our dear friend and colleague Pierre-Francois Van de Moortele who recently passed and was a major contributor to this and many works of our group. This research was funded by NIH grants U01 EB025144, P41 EB027061, P30 NS076408, S10 RR029672, and R01 EB024536.

## SUPPORTING INFORMATION

### RF COIL DESIGN STRATEGIES for IMPROVING SNR at the ULTRAHIGH MAGNETIC FIELD of 10.5 TESLA

#### 1. 10.5T 16-channel Tx/Rx Array

The 16-channel SD array was laid out in two rows along the z-axis, with each row consisting of eight azimuthally distributed elements, placed at a diameter of 28 cm. This two-row design, which we had employed previously (1,2) and was subsequently pursued by others (e.g. (3–5)), was motivated by previous work demonstrating that it provides an optimal geometry for parallel transmit (pTx) pulse design for minimizing SAR (6). Each coil element was a 9.15 x 9.15 cm square (conductor center-to-center), with a trace width of 7.5 mm. The two rows of eight coil elements (Figure 1C) were offset from one another by 22.5 degrees azimuthally with an 11 mm gap between coil conductors in each row as well as between rows. All conductors of the 16-channel SD array were made from 0.5 oz (18 µm) copper with a silver immersion finish on a flexible polyimide substrate with a thickness of 0.1 mm. Each of the square-shaped SD array coil elements (Figure 1B) was divided into four segments by cuts at each of the four corners. Multilayer ceramic chip capacitors (100C Series, American Technical Ceramics, Huntington Station, NY, USA) were located at three of the four corners, while the fourth corner hosted a pair of 1-12 pF variable capacitors (V9000, Knowles-Voltronics Corp, Cazenovia, NY, USA) that allowed for the tuning and matching each coil element.

A coil base was also designed to fixate the coil assembly to the contours of the patient table. All coil housings and base components were designed and 3D printed in-house by an additive manufacturing process using a polyethylene terephthalate glycol (PETG) filament (Atomic Filament, Kendallville, IN) on a Fusion3 F410 (Fusion3, Greensboro, North Carolina) printer; no surface finishes were applied to the coil housings.

#### 2. 10.5T Integrated MR System Interface

The T/R switches were fabricated using all non-magnetic components on 1.6 mm thick FR-4 printed circuit board material with 3.5 mil trace thickness (1 oz copper weight) and individually optimized for 447 MHz operation. Non-magnetic MCX coaxial contacts (Cinch Connectivity Solutions, Waseca, MN, USA) were placed at each RF port of the T/R switch to provide modularity to the device assembly to allow for optimization, measurement, and ease of service if it were necessary. Supporting Information Figure S1 shows the T/R switch schematic. A variety of the ceramic chip capacitors (100B series, American Technical Ceramics, Huntington Station, NY, USA) along with wire wound inductors (1512SP & 1812SMS series, Coilcraft Inc., Cary, IL, USA) were used for the assembly and optimization of each T/R switch. All of the detuning signal paths used to transition the T/R switch from transit to receive mode by way of both series and shunt PIN diodes (MA4P7470-1072T, MACOM, Lowell, MA, USA) were filtered via a combination of ceramic-core RF chokes (1008CS series, Coilcraft Inc., Cary, IL, USA) and ceramic chip capacitors (0603 Series, Knowles Corp, Itasca, IL, USA). The switching speed observed was ∼2.5 µs when transitioning into the transmit state and ∼8.5 µs when transitioning into the receive state while using the PIN switching driver signals provided by the MR system. A low input impedance preamplifier (WMM447P, WanTcom, Chanhassen, MN, USA) was integrated into each T/R switch assembly (Figure 2C), which provided adequate gain of the received signal back to the MR system. Each T/R switch was connected to its respective SD coil element with a quarter-wavelength coaxial cable (SFT-316, Times Microwave Systems, Wallingford, CT, USA). This λ/4 length cable along with circuitry integrated into the T/R switch facilitated the impedance transformation necessary to achieve preamplifier decoupling (7) of the SD elements during the acquisition period in order to reduce coupling to the receive-only array. Resonant cable traps in the form of an electrically-shortened “bazooka” balun (8) were added on the system side of each T/R switch to mitigate MR system cable interactions, while sleeve-type ”bazooka” baluns were added to each SD coil element feed cable to reduce common mode currents from the coil elements, as well as reduce cable-to-coil interaction between rows. Additional resonant cable traps were added to the coaxial feed cables of the superior elements to attenuate any feed cable interactions with its associated coil element. The individual transmit cables within the two 8-channel parallel transmit (pTx) MR system plug assemblies were cut to length in order to reach each respective T/R switch location within the coil housing.

**SUPPLEMENTARY FIGURE S1:**
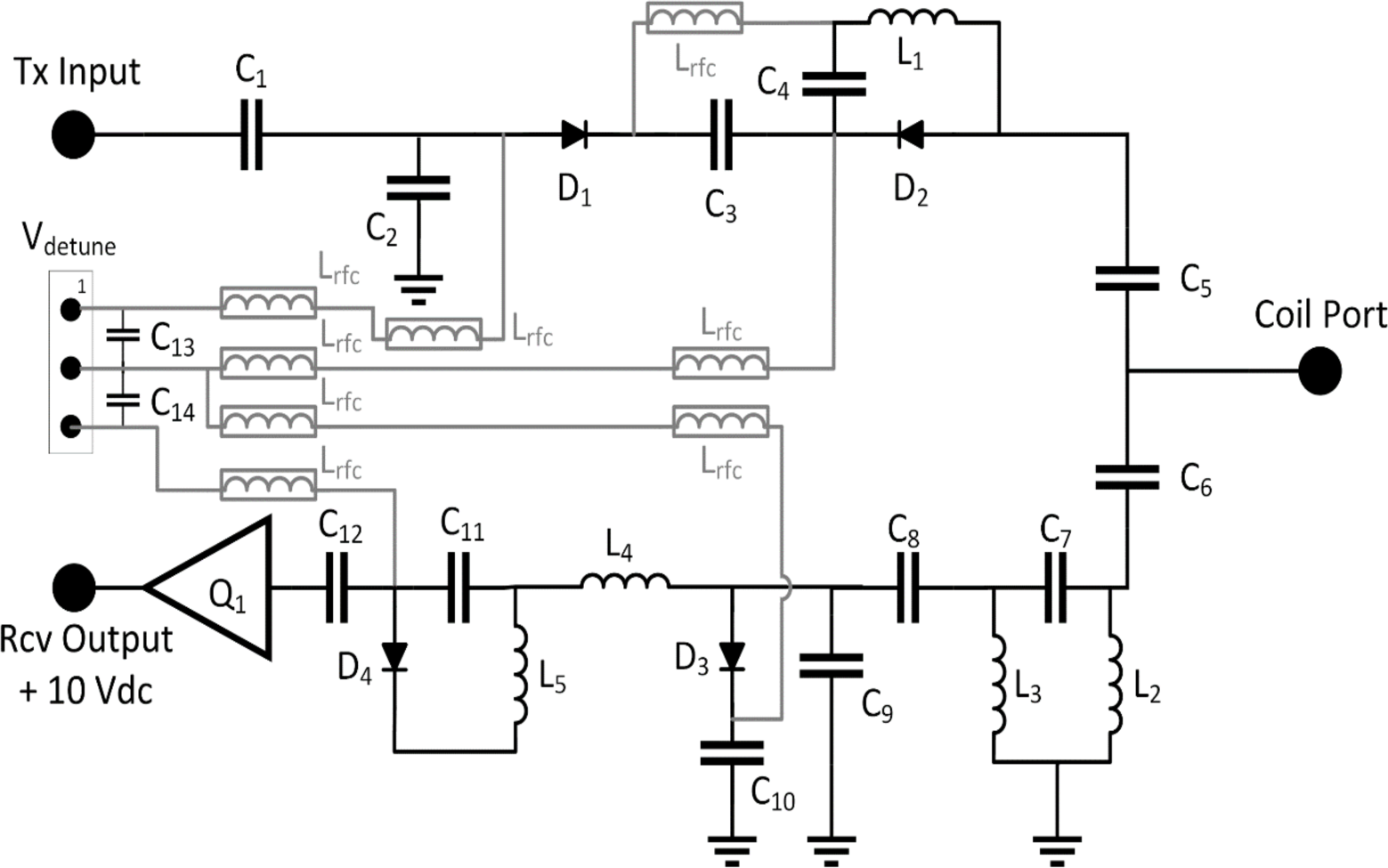
Schematic of an integrated T/R switch circuit. Light grey traces and components indicate V_dc_ paths, while black traces indicate RF paths.

#### 3. 10.5T 64-channel Receive-Only Array

Each receive-only coil element was constructed using 18 AWG silver-plated copper wire (MWS Wire Industries, Oxnard, CA, USA) formed directly onto the coil housing (Figure 3B-C). The representative schematic (Figure 3A) shows ceramic chip capacitors (C_1_, C_2_, C_dt_) (1111 Series, Knowles Corp, Itasca, IL, USA) used to distribute the tuning capacitance around the loop, with the fine tuning done with a variable capacitor (C_t_) (SGC3 series, Sprague Goodman Electronics Inc., Westbury, NY, USA). The feed circuit location included a pair of variable capacitors (C_m1_, C_m2_) (Thin-Trim series, Johanson Manufacturing Corp., Booton, NJ, USA) for coil matching, active and passive detuning circuitry using cross-pair diodes (D_dt_) (UMX9989AP, Microsemi, Lowell, MA, USA) with inductor (L_dt_) (Square Air Core series, Coilcraft Inc., Cary, IL, USA) in parallel with capacitor C_dt_, and preamplifier protection circuitry with a fast switching PIN diode (D_p_) (MA4P1250NM-1072T, MACOM, Lowell, MA, USA) and series inductor (L_p_) (Square Air Core series, Coilcraft Inc., Cary, IL, USA) in parallel with a multilayer ceramic chip capacitor (C_p_) (1111 Series, Knowles Corp, Itasca, IL, USA). All coil elements were tuned and matched to the lightbulb-shaped phantom loading condition described above.

Each receive-only coil element was connected to a low input impedance preamplifier (WMA447D, WanTcom, Channhassen, MN, USA) located at the superior end of the coil assembly by a length of low-loss semi-rigid coaxial cable (UT-047C-TP-LL, Micro-Coax Inc., Pottstown, PA, USA). Each of these feed cables included a resonant cable trap to minimize interaction with the 16-channel SD array. Variable capacitors (SGC3 series, Sprague Goodman Electronics Inc., Westbury, NY, USA) located at the preamplifier input were used to optimize preamplifier decoupling during acquisition.

The 7T 64Rx array was tuned and matched to better than -15dB at 297 MHz using the lightbulb phantom. Similar to the 10.5T 64Rx array, bench measurements of two 25×50 mm2 7T elements demonstrated -34.0 dB coupling while overlapped, and ∼-7 dB coupling while non-overlapped, relying on the use of preamplifier decoupling to further isolate the coils. Similar results were observed for the 50×50 mm2 elements. QU to QL ratios ranged from 5:1 for heavier loading conditions to 2:1 for lighter loading conditions. Typical active PIN detuning of ∼30 dB was achieved with the 7T 64Rx.

#### 4. Safety Validation

##### a) EM Modeling

The EM modeling was performed following the workflow detailed in previous work (9). The 16-channel Tx coil was modeled using the exact layout shown in Figure 1, implemented as perfect electrical conductor (PEC) strips. The coil former was modeled as PETG with electrical properties of Ɛr = 1.7 and σ = 4.2×10-5 S/m. As previously mentioned, the electrical properties of the gel (i.e., PVP + AGAR + NaCl solution) in the lightbulb phantom employed in the experimental measurements as well as the EM simulations, were measured as σ = 0.65 S/m and Ɛr = 47.2. The EM model of the Tx coil consisted of 96 ports, six ports per loop, with five ports representing tuning/matching capacitors and one port representing the RF feed. All EM simulations were performed in a frequency domain EM simulator (HFSS, Ansys, Canonsburg, PA) with the finite element method (FEM) solver. An iterative adaptive mesh refinement tool was used to refine the non-uniform tetrahedral meshes to resolve the model. This mesh refinement technique was able to produce approximately 1 million tetrahedral meshes with successive scattering parameters (S-parameters) changing less than 10^-3^ at 447 MHz. The resulting S-parameters were imported into a circuit simulator (AWR Corp., El Segundo, CA) to optimize the value of the lumped elements and to minimize the root mean square error (RMSE) between simulated and measured S-parameters. Optimized lumped element values were then incorporated into the EM model to generate B_1_^+^ maps, which were compared to experimentally measured B_1_^+^ maps. EM modeling, circuit co-simulation, and post-processing were performed on a workstation with two 8-core Intel(R) Xeon(R) W-2245 processors with a 3.9 GHz clock rate and 256 GB RAM.

**SUPPLEMENTARY FIGURE S2:**
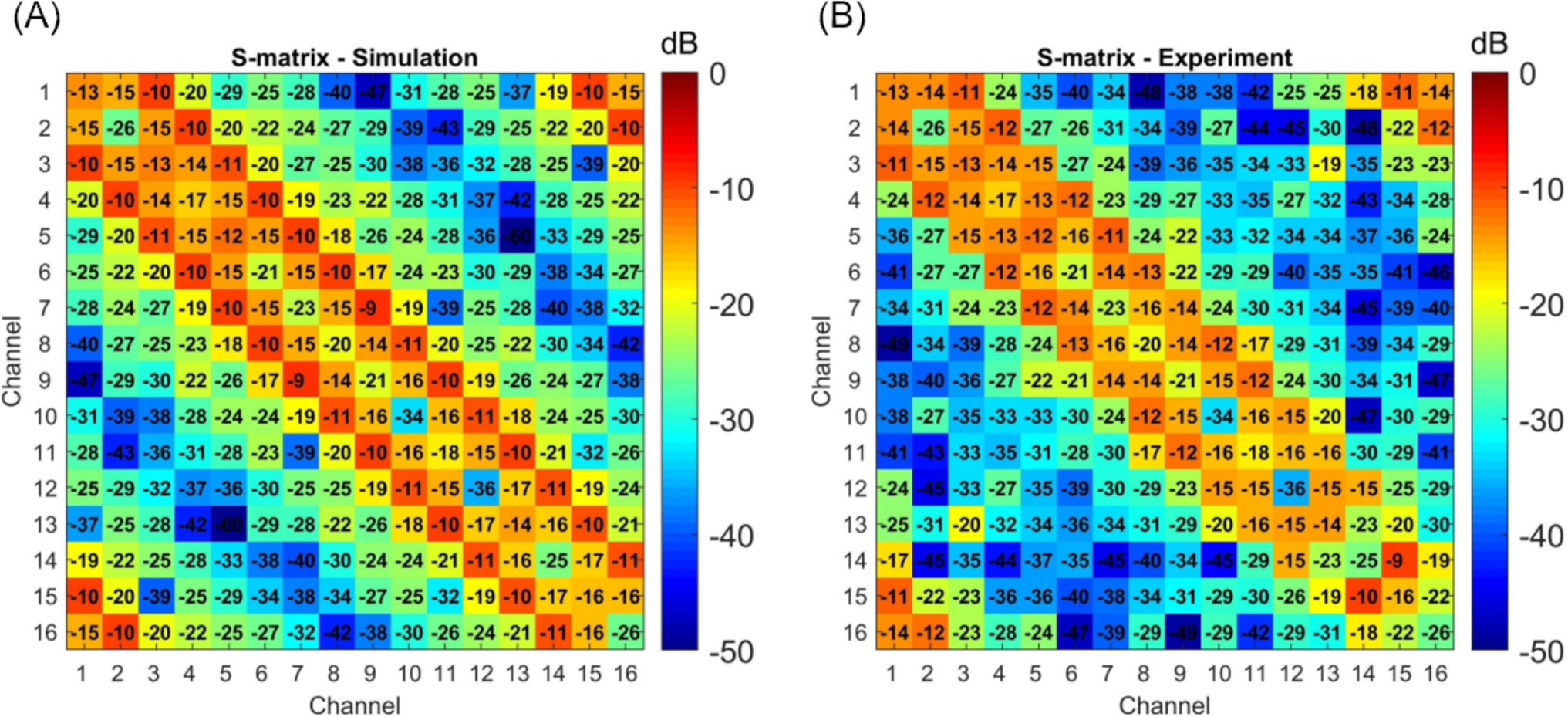
(A) Simulated S-parameter matrix of the 16-channel SD transmitter alone, and (B) experimental S-parameter matrix of the 16-channel SD transmitter (SD_i_) when used in conjunction with the 64-channel receive-only insert.

**SUPPLEMENTARY FIGURE S3:**
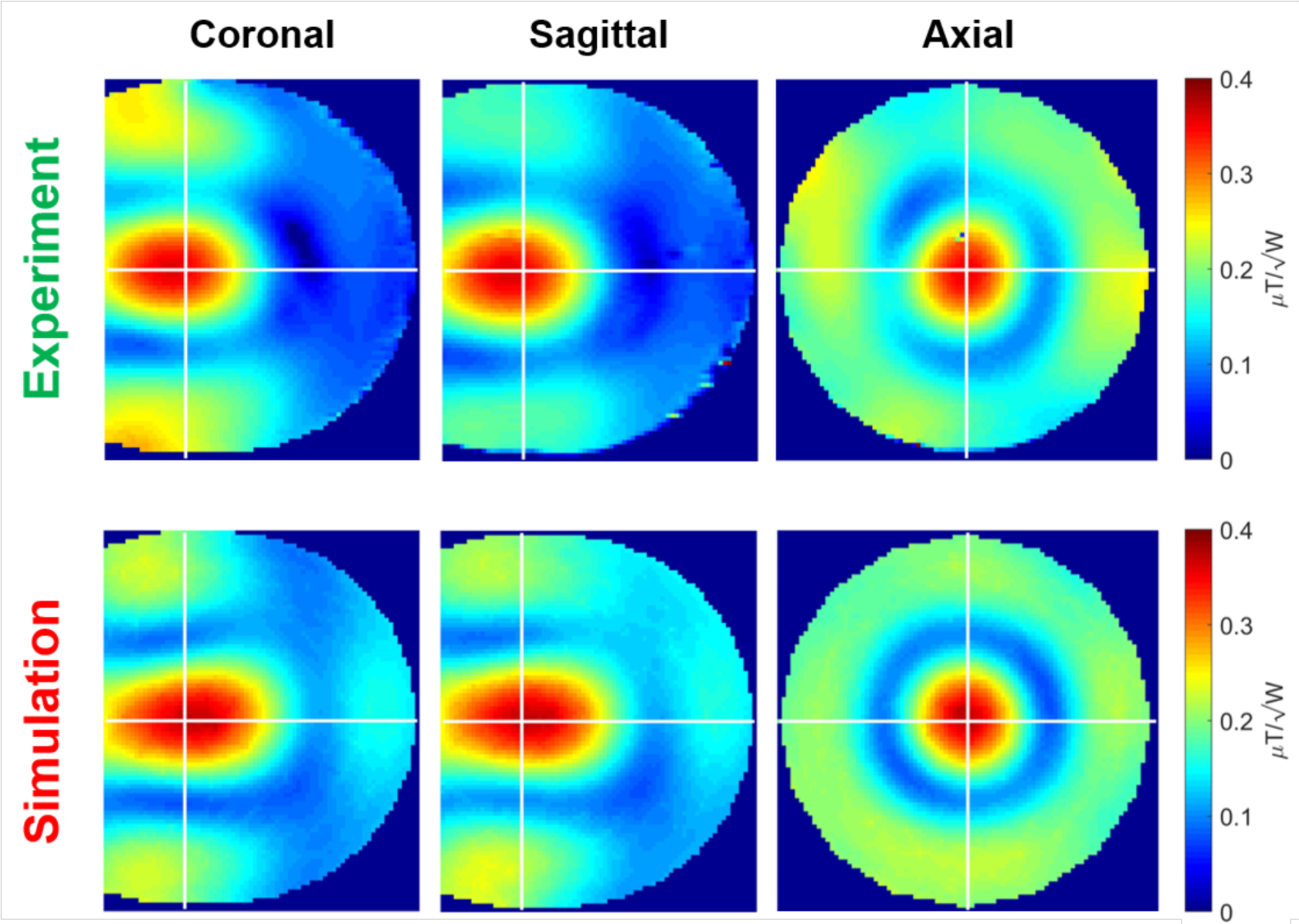
Simulated and experimental B_1_^+^ power efficiency maps corresponding to a CP-like excitation pattern are shown in the 16Tx/80Rx array that uses the 16-ch SD array (SD_i_) for transmission. The CP-like solution was obtained by RF shimming, targeting and achieving the highest power efficiency at the center of the phantom. Simulated maps were calculated using the voltages reported by the MR scanner. The results are used to calculate an NRMSE, quantifying the error between simulated and measured B_1_^+^ data.

##### b) Safety Factor Calculation

A safety factor (SF) is critical for accounting for uncertainties in peak spatial 10g-averaged SAR (psSAR10g) induced in a subject (*^e_SAR_^*). The SF was used to scale up the predicted psSAR10g and scale down the input power level to ensure patient safety.

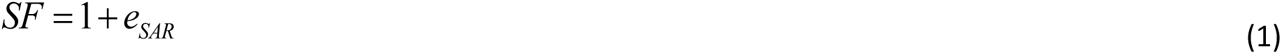

As previously proposed (10,11) three sources of uncertainty were considered for *^e_SAR_^* in the SAR analysis, namely, power monitoring uncertainty (*^e_PM_^*), inter-subject variability (*^e_ISV_^*), and EM modeling uncertainty (*^e_EMM_^*). These three were then combined with a sum-of-squares approach to calculate the uncertainty in the psSAR10g prediction (11):

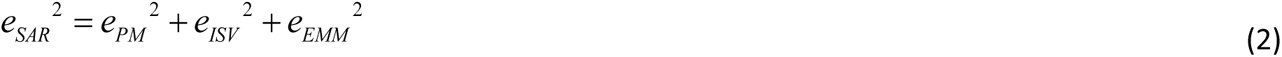

Previously proposed *e*_*ISV*_ value (12) and the *e*_*PM*_ reported by the MR system manufacturer (i.e., power measurement uncertainty of the directional couplers) were used in the *e*_*SAR*_ calculation. To quantify the *e*_*EMM*_, the error between simulated and measured B1+ maps was propagated to calculate the uncertainty in the psSAR10g prediction, as outlined in detail in previous work by Sadeghi-Tarakameh et al (13).

**SUPPLEMENTARY FIGURE S4:**
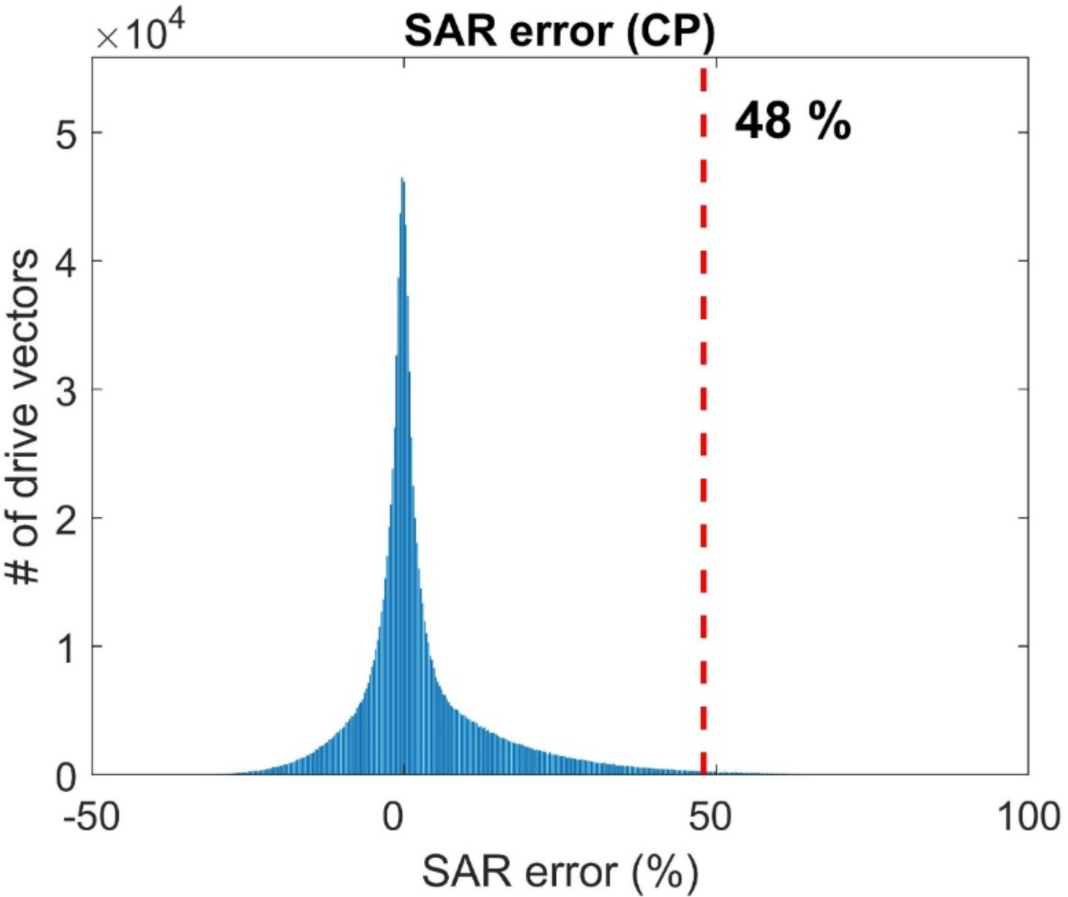
Histogram of the SAR-error is obtained from Monte-Carlo simulations. The plot shows the distribution of SAR error obtained from different RF excitation scenarios satisfying the condition B_1_^+^ NRMSE ≤ 25%. For 99.9% of all RF excitation scenarios, psSAR_10g_ deviated less than 48% from the psSAR10g of the CP-like mode.

##### c) Power Limit Calculation

The EM model of the 16 channel Tx coil (obtained in phase one, with optimal lumped elements) was used to simulate the electric field distribution in a realistic human head model (The Ansys Human Body Model, Ansys, Canonsburg, PA) and to calculate the Q-matrices (14). Similar to the EM simulation in the first phase, an iterative adaptive mesh refinement tool was used, and approximately 2 million tetrahedral meshes were generated. Then the Q-matrices were scaled up by the safety factor and compressed using the virtual observation point (VOP) technique (15) with a 10% overestimation factor. During in vivo scans, these VOPs were used to evaluate the psSAR10g and to ensure patient safety for different RF shimming scenarios.

#### 5. Data Acquisition Processing: Functional Imaging

For the visual stimulation we used face stimuli; we collected 5 runs of a face perception task where participants were instructed to report whether they perceived a face. We manipulated the phase coherence of grey scale images (from 0% to 40% in steps of 10%), effectively manipulating the visibility of the face stimuli. Stimuli subtended approximately 9 degrees of visual angle. Each image was cropped to remove external features using an ellipse. Ellipses spanned the full vertical extent of each face and 80% of the external extent. The edge of the ellipse was smoothed by convolution with an average filter (using "fspecial" with "average" option in MatLab (Mathworks, Natick, Massachusetts, USA). This procedure was implemented in order to prevent participants from performing simple edge detection using the hard edges typically present in face images. We normalized the amplitude spectrum of the face images by ensuring that the rotational average amplitudes for a given spatial frequency were equated across stimuli while preserving the amplitude distribution across orientations. The standard deviation of pixel intensity was also kept constant across stimuli. These images were used in a pseudo-block design with 10 stimuli presented in a 1.2s on/1.2s off sequence for a total block duration of 24 seconds with an inter-bloc interval of 24 seconds. During the “off” and the inter-bloc intervals, a grey background was shown to the participants. A fixation cross was kept onscreen throughout the visual stimulation. For each run, 8 blocks were shown, with an additional 24 seconds "off" period at the beginning of the scan producing a total run length of 408s. Participants were instructed to maintain fixation on the centered crosshair throughout the entire run.

Functional pre-processing included 3D rigid body motion correction (where each volume of each run was realigned to the first volume of the first run using sinc interpolation) and linear detrending, performed by regressing out low-drifts, up to 3rd order discrete cosine transform from the motion-corrected time series. Subsequently, standard general linear model (GLM) was performed to estimate BOLD-evoked response amplitudes using ordinary least squares minimization. GLM design matrices were generated by convolving of a double gamma with a "box car" function, the latter representing the onsets and offsets of the stimuli. GLM analyses were performed on each run independently.

#### 6. RESULTS

##### a) Noise Correlation (Suppl. Fig. S5)

**SUPPLEMENTARY FIGURE S5:**
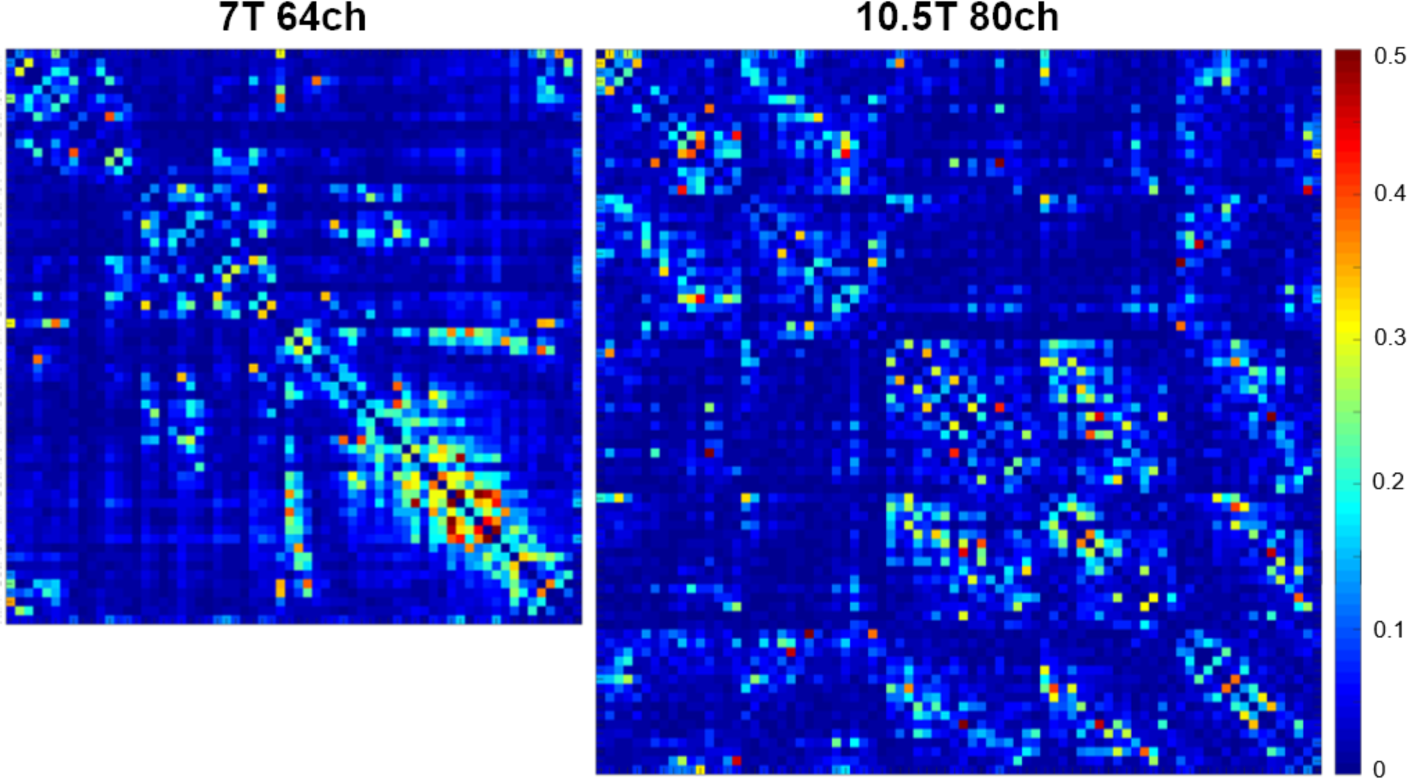
(A) Experimentally measured 80-channel noise correlation matrix of the 16Tx/80Rx array while loaded with a human head (80Rx channels are composed of the 64-channel receive-only array paired with the 16-channel SD_i_ transceiver array). The average noise correlation for 80-channels was 4.96% with a phantom load and 4.46% with a human subject.

##### b) SNR Improvement with the Integrated System Interface (Suppl. Fig. S6)

**SUPPLEMENTARY FIGURE S6:**
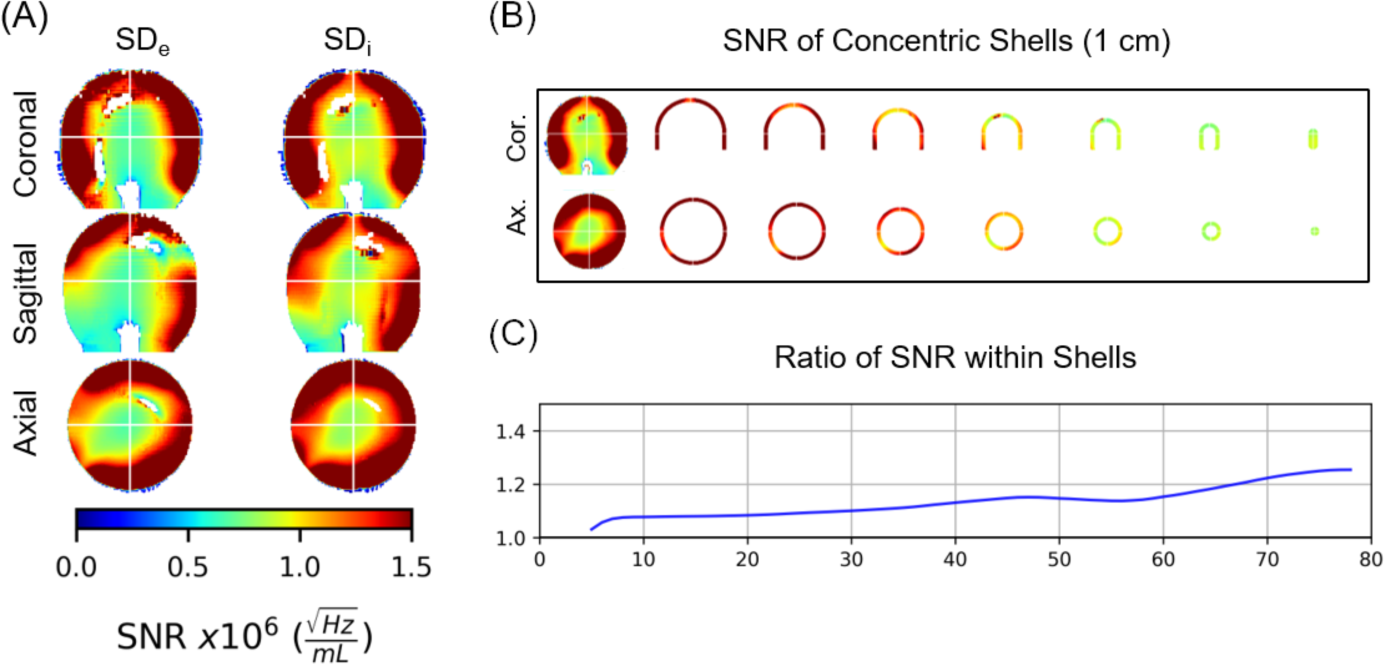
Demonstration of SNR improvements within a phantom achieved utilizing the integrated MR system interface (SD_i_) vs. the external interface (SD_e_). (A) SNR plots across the central slice in three imaging planes. (B) SNR of the SD_i_ array shown in 1cm thick concentric shells. (C) The ratio of SNR (SD_i_/SD_e_) as a function of depth resolved by 1 cm thick concentric shells.

##### c) SNR Improvement with addition of the Transmitter as a Receiver (Suppl. Fig. S7)

**SUPPLEMENTARY FIGURE S7:**
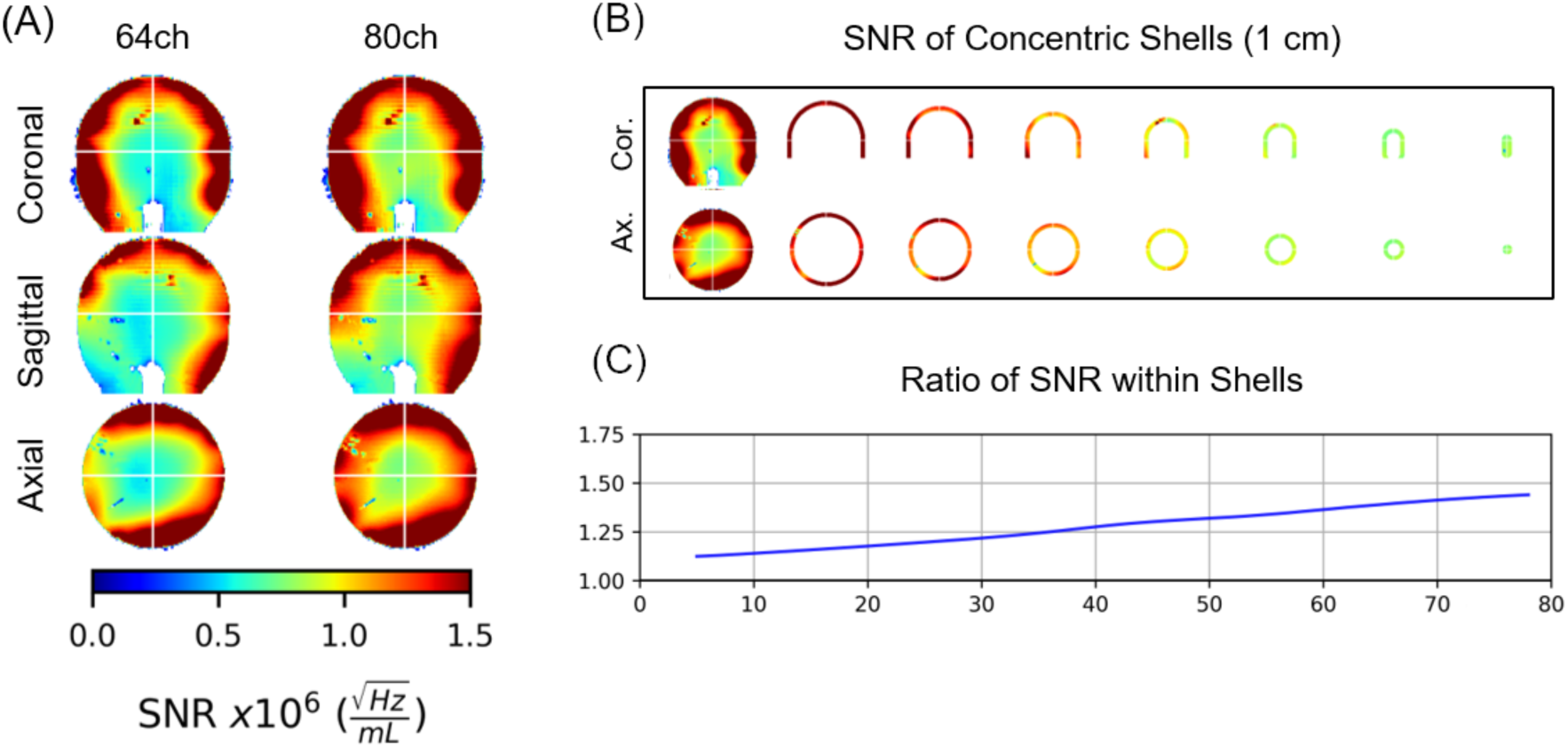
SNR gains at 10.5T when utilizing a 64-channel receive-only array together with a 16-channel SD transmitter array (SDi) operating as a transceiver for a combined 80-channel receive array. (A) SNR maps across in three orthogonal central slices for 64-channel receive only array and the combined 80-channel receive array; right most column in (A) illustrates the ratio of the two. (B) SNR of the 80ch array shown in 1cm thick concentric shells in the same three orthogonal planes shown in (A). (C) The ratio of SNR (80ch/64ch) averaged in the 1 cm thick concentric shells as a function of depth

#### 7. DISCUSSION

##### a. New SD TRANSMITTER vs the INITIAL INDUCTIVELY COUPLE TRANSMITTER

**SUPPLEMENTARY FIGURE S8:**
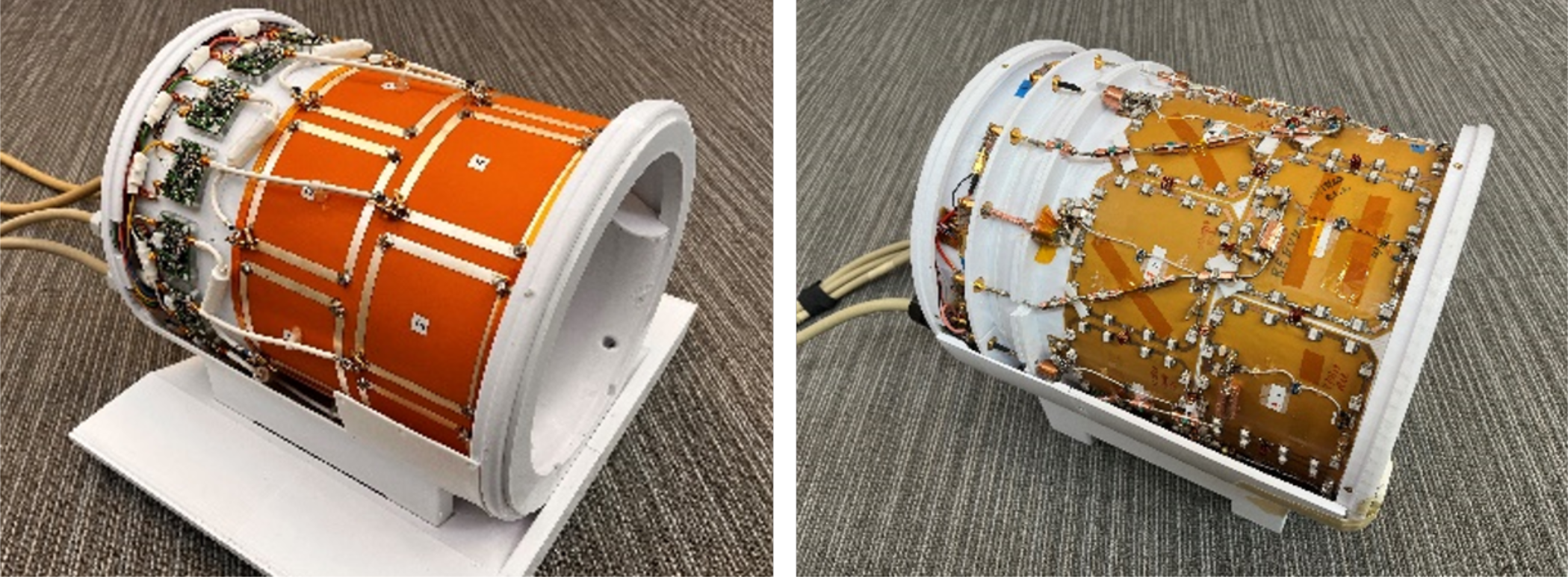
The simplicity of the current self-decoupled 16Tx/Rx SD array (Left), can be appreciated when compared to our initial inductively-decoupled 16Tx array (Left) which used conventional loops segmented by multiple capacitors. The two coils arrays had similar layouts utilizing two rows of 8 loops azimuthally distributed; however, the minimal component count in the SD design led to a more streamlined simulation and safety validation process.

##### b. <u>SUPLEMANTARY FIGURE S9:</u> ADDITIONAL 10.5T ANATOMICAL IMAGES and/or EXPANDED VERSIONS of the 10.5T IMAGES PRESENTED in the MAIN BODY of the PAPER in FIGURE 9

**SUPPLEMENTARY FIGURE S9.1:**
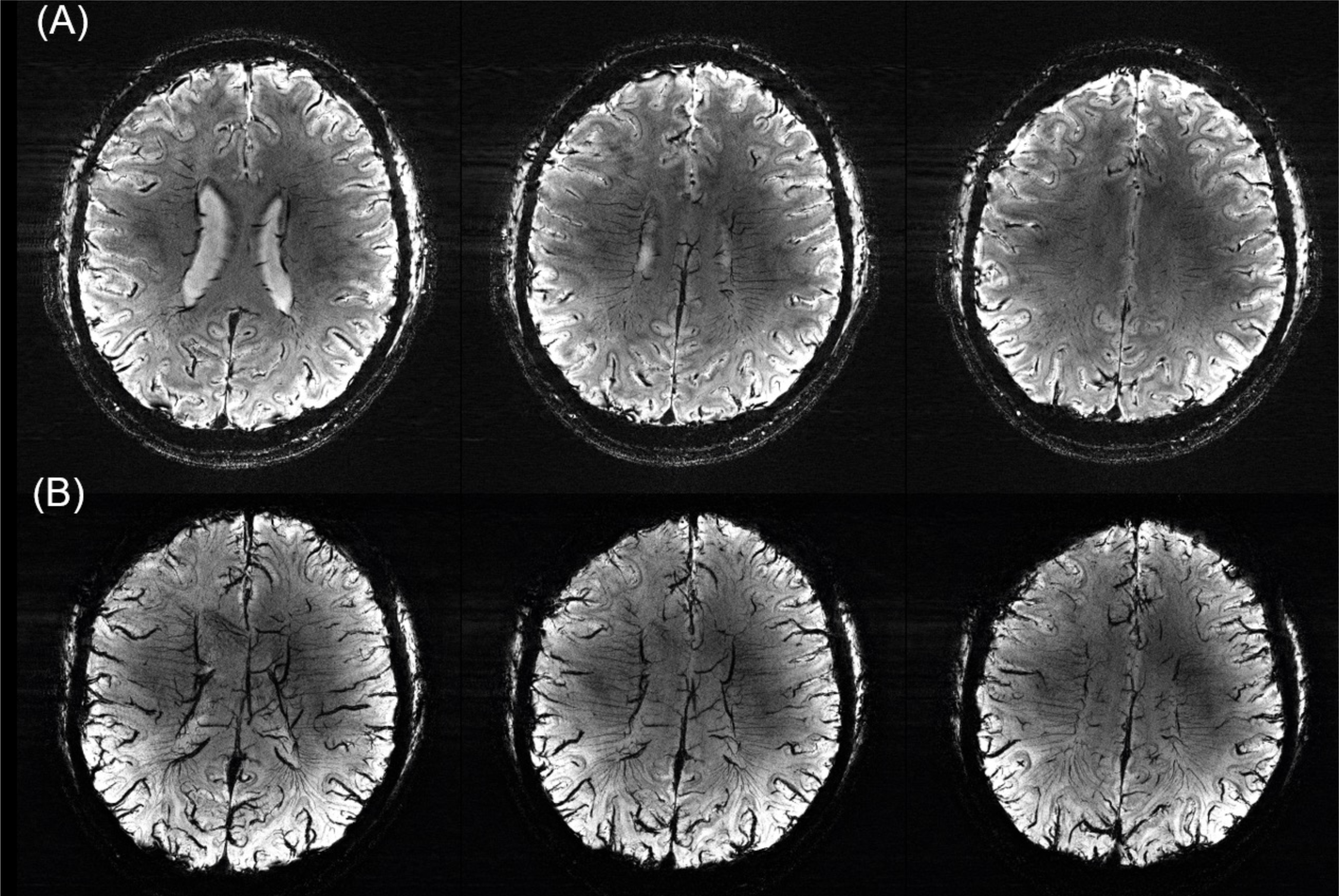
(A) Example of susceptibility weighted imaging (SWI) for three representative single slices (1.3mm thick), and (B) the minimum intensity projection (SWI mIP) for a 10.4 mm thick slab derived from the SWI data of the human brain at 10.5T acquired using the 16Tx/80Rx head RF coil array in 36 contiguous axial-coronal oblique slices at 0.21 x 2.1 mm2 in-plane and 1.3 mm slice resolution using a 3D GRE sequence. Other relevant imaging parameters were: TR = 35 ms, TE = 18 ms, FOV = 215 (read) x 188 (phase) x 46.8 (slice) mm3, in-plane acceleration factor along phase encode direction (i.e. in-plane reduction factor or in Siemens terminology iPAT factor) = 5, nominal flip angle = 14°, and bandwidth = 315 Hz/pixel.

**SUPPLEMENTARY FIGURE S9.2:**
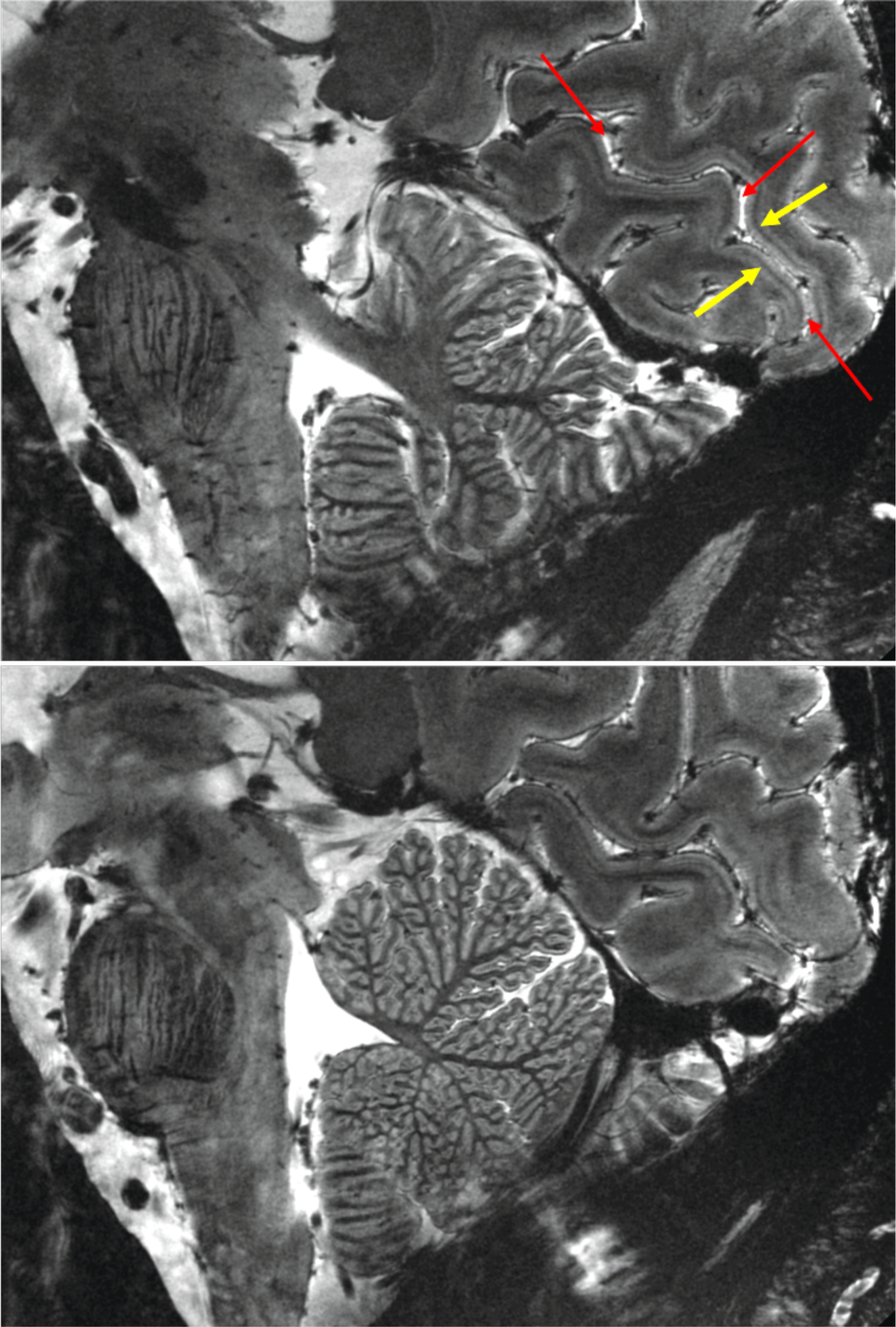
The sagittal GRE Images from Figure 9 in the main body of the paper-expanded and some structures labeled (colored arrows): The calcarine fissure is identified with the RED ARROWS (upper panel) at each end and the middle. The upper and lower banks of the calcarine fissure comprises the primary visual cortex or V1 identified with a unique a white matter stria (stria or line of Gennari) identified by the YELLOW ARROWS. 0.2×0.2 MM in plane resolution and 1 mm slice thickness; ∼5 min data acquisition; factor of 2 undersampling along the phase encode direction.

**SUPPLEMENTARY FIGURE S9.3:**
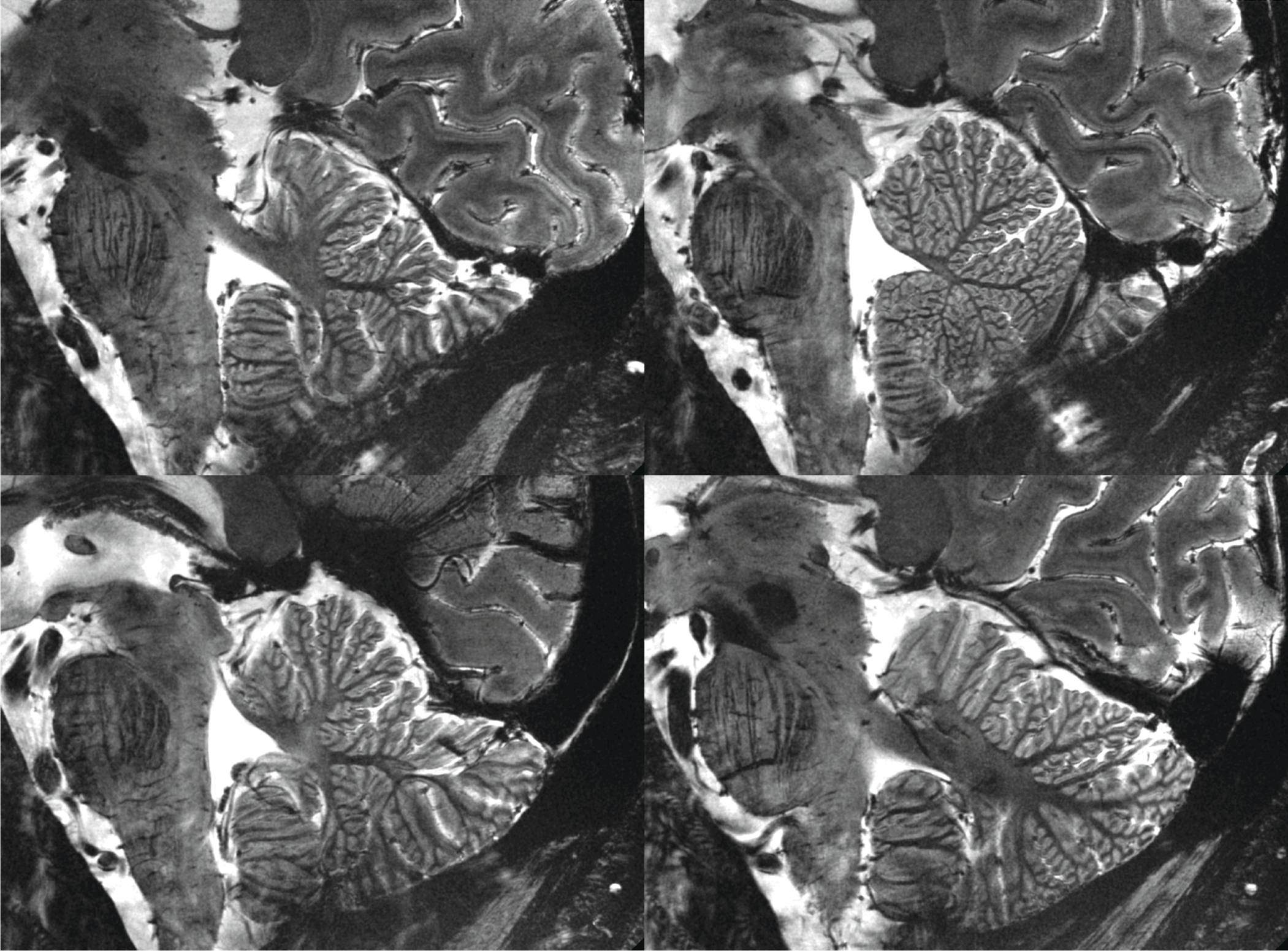
The sagittal GRE Images from Figure 9 in the main body of the paper and Supplemental Figure S9.2, showing 4 slices.

**SUPPLEMENTARY FIGURE S9.4:**
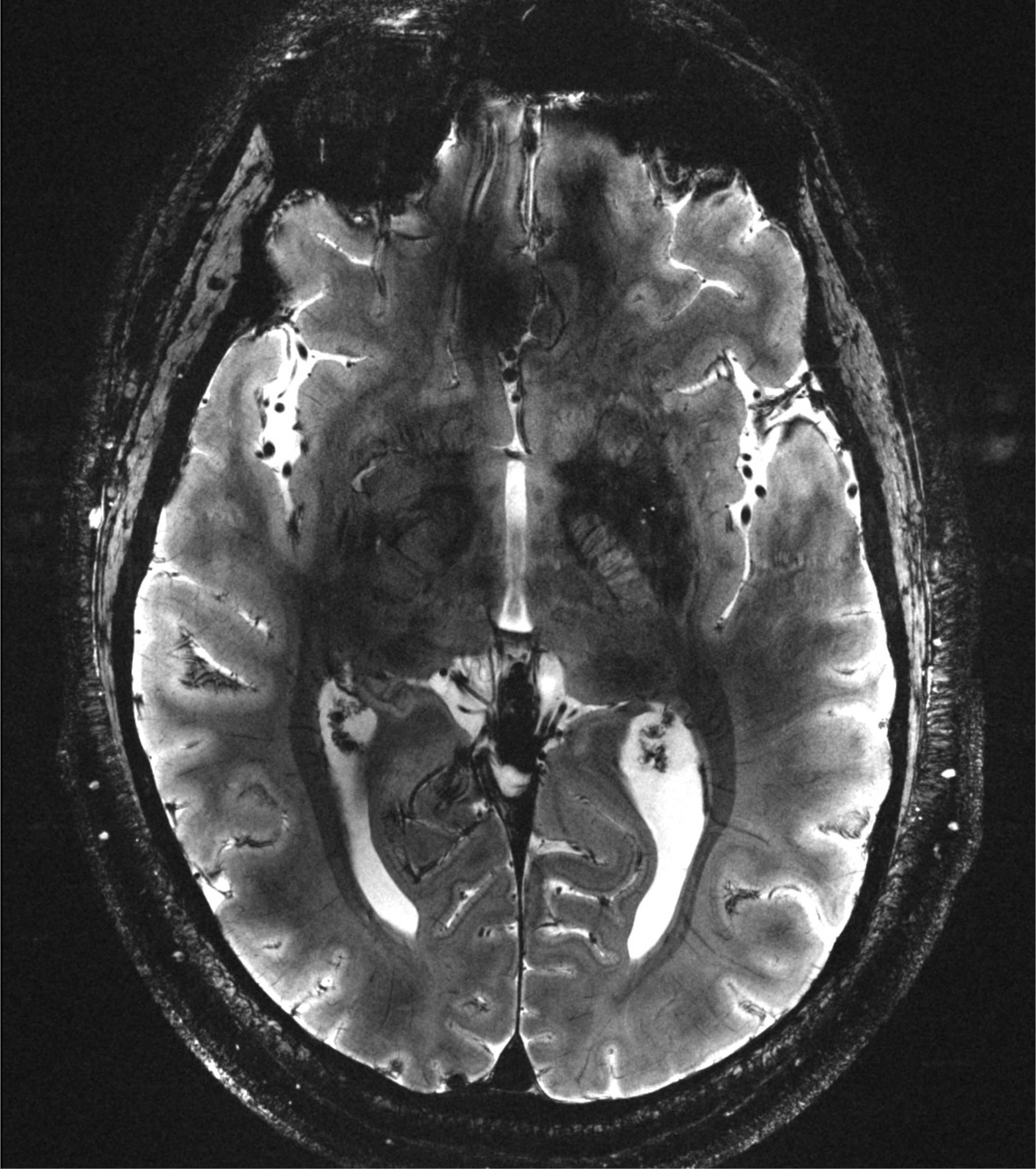
One of the two axial slices of the Gradient recalled echo images shown in Figure 9 in the main body of the paper, presented in an enlarged form to appreciate the details of the image. ∼5 min of data acquisition.0.2×0.2 mm in plane resolution and 1 mm slice thickness.

**SUPPLEMENTARY FIGURE S9.5:**
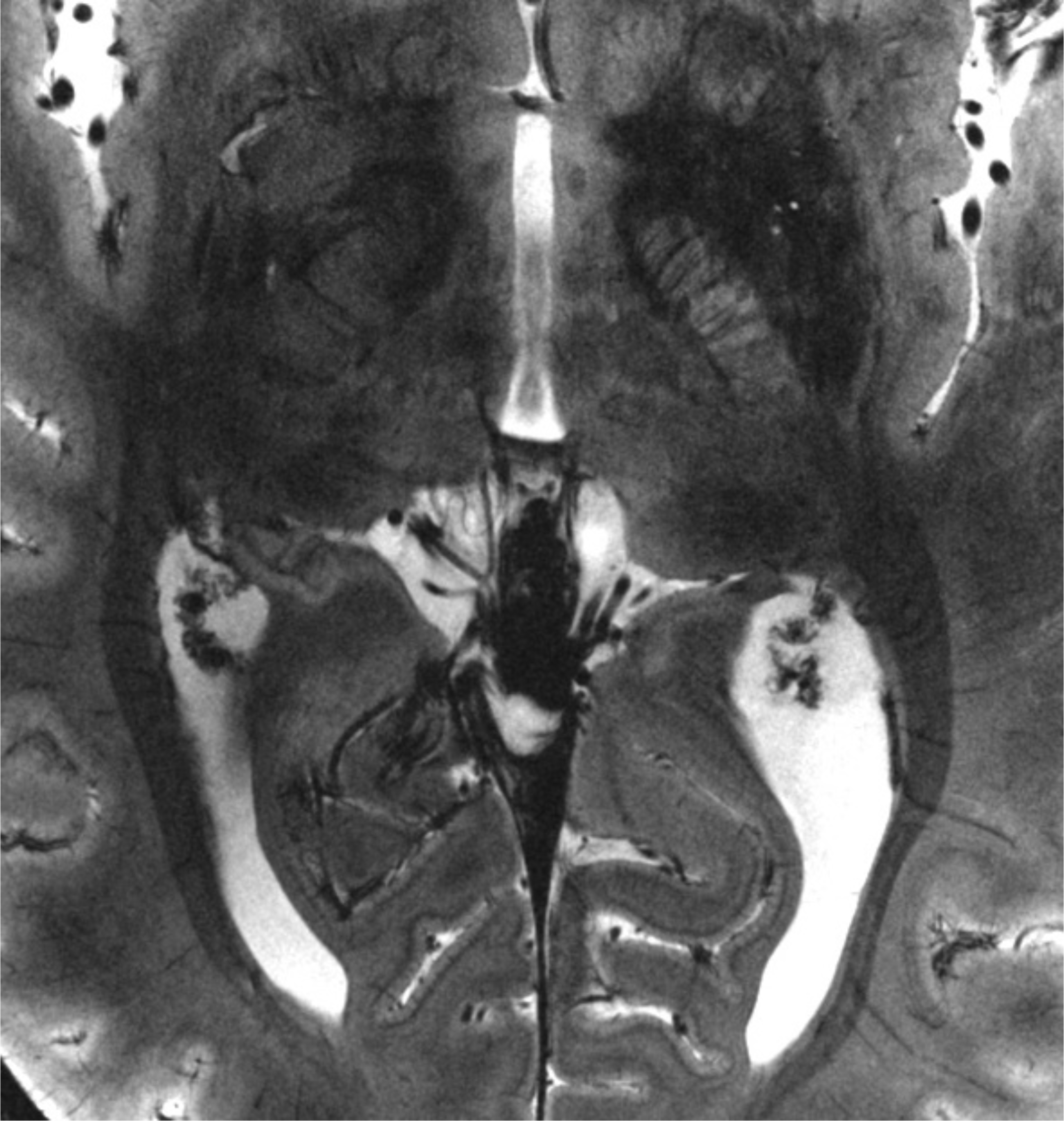
An expanded view of the middle section of the slice shown in Supplemental Figure S9.4. Optic radiation are the dark bundles, one on each hemisphere, that run along the ventricles, which seen as bright white regions. Intracortical veins that run perpendicular to the cortex are seen in many places.

## References

1. Ugurbil K. Imaging at ultrahigh magnetic fields: History, challenges, and solutions. Neuroimage 2018;168:7–32. doi:10.1016/j.neuroimage.2017.07.007

2. Ugurbil K. Magnetic resonance imaging at ultrahigh fields. IEEE Trans Biomed Eng 2014;61(5):1364–1379. doi:10.1109/TBME.2014.2313619

3. Dumoulin SO, Fracasso A, van der Zwaag W, Siero JCW, Petridou N. Ultra-high field MRI: Advancing systems neuroscience towards mesoscopic human brain function. Neuroimage 2018;168:345–357. doi:10.1016/j.neuroimage.2017.01.028

4. De Martino F, Yacoub E, Kemper V, Moerel M, Uludag K, De Weerd P, Ugurbil K, Goebel R, Formisano E. The impact of ultra-high field MRI on cognitive and computational neuroimaging. Neuroimage 2018;168:366–382. doi:10.1016/j.neuroimage.2017.03.060

5. Yacoub E, Harel N, Ugurbil K. High-field fMRI unveils orientation columns in humans. Proc Natl Acad Sci U S A 2008;105(30):10607–10612. doi:10.1073/pnas.0804110105

6. Huber L, Finn ES, Chai Y, Goebel R, Stirnberg R, Stocker T, Marrett S, Uludag K, Kim SG, Han S, Bandettini PA, Poser BA. Layer-dependent functional connectivity methods. Prog Neurobiol 2021;207:101835. doi:10.1016/j.pneurobio.2020.101835

7. Santoro R, Moerel M, De Martino F, Goebel R, Ugurbil K, Yacoub E, Formisano E. Encoding of natural sounds at multiple spectral and temporal resolutions in the human auditory cortex. PLoS Comput Biol 2014;10(1):e1003412. doi:10.1371/journal.pcbi.1003412

8. Cai Y, Hofstetter S, van der Zwaag W, Zuiderbaan W, Dumoulin SO. Individualized cognitive neuroscience needs 7T: Comparing numerosity maps at 3T and 7T MRI. Neuroimage 2021;237:118184. doi:10.1016/j.neuroimage.2021.118184

9. Vu AT, Jamison K, Glasser MF, Smith SM, Coalson T, Moeller S, Auerbach EJ, Ugurbil K, Yacoub E. Tradeoffs in pushing the spatial resolution of fMRI for the 7T Human Connectome Project. Neuroimage 2017;154:23–32. doi:10.1016/j.neuroimage.2016.11.049

10. Ozutemiz C, White M, Elvendahl W, Eryaman Y, Marjanska M, Metzger GJ, Patriat R, Kulesa J, Harel N, Watanabe Y, Grant A, Genovese G, Cayci Z. Use of a Commercial 7-T MRI Scanner for Clinical Brain Imaging: Indications, Protocols, Challenges, and Solutions-A Single-Center Experience. AJR Am J Roentgenol 2023;221(6):788–804. doi:10.2214/AJR.23.29342

11. Ladd ME, Bachert P, Meyerspeer M, Moser E, Nagel AM, Norris DG, Schmitter S, Speck O, Straub S, Zaiss M. Pros and cons of ultra-high-field MRI/MRS for human application. Prog Nucl Magn Reson Spectrosc 2018;109:1–50. doi:10.1016/j.pnmrs.2018.06.001

12. Ugurbil K, Van de Moortele PF, Grant A, Auerbach EJ, Erturk A, Lagore R, Ellermann JM, He X, Adriany G, Metzger GJ. Progress in Imaging the Human Torso at the Ultrahigh Fields of 7 and 10.5 T. Magn Reson Imaging Clin N Am 2021;29(1):e1–e19. doi:10.1016/j.mric.2020.10.001

13. Sadeghi-Tarakameh A, DelaBarre L, Lagore RL, Torrado-Carvajal A, Wu X, Grant A, Adriany G, Metzger GJ, Van de Moortele PF, Ugurbil K, Atalar E, Eryaman Y. In vivo human head MRI at 10.5T: A radiofrequency safety study and preliminary imaging results. Magn Reson Med 2020;84(1):484–496. doi:10.1002/mrm.28093

14. He X, Erturk MA, Grant A, Wu X, Lagore RL, DelaBarre L, Eryaman Y, Adriany G, Auerbach EJ, Van de Moortele PF, Ugurbil K, Metzger GJ. First in-vivo human imaging at 10.5T: Imaging the body at 447 MHz. Magn Reson Med 2020;84(1):289–303. doi:10.1002/mrm.28131

15. Grant A, Metzger GJ, Van de Moortele PF, Adriany G, Olman C, Zhang L, Koopermeiners J, Eryaman Y, Koeritzer M, Adams ME, Henry TR, Ugurbil K. 10.5 T MRI static field effects on human cognitive, vestibular, and physiological function. Magn Reson Imaging 2020;73:163–176. doi:10.1016/j.mri.2020.08.004

16. Schmidt S, Erturk MA, He X, Haluptzok T, Eryaman Y, Metzger GJ. Improved (1) H body imaging at 10.5 T: Validation and VOP-enabled imaging in vivo with a 16-channel transceiver dipole array. Magn Reson Med 2024;91(2):513–529. doi:10.1002/mrm.29866

17. Le Ster C, Grant A, Van de Moortele PF, Monreal-Madrigal A, Adriany G, Vignaud A, Mauconduit F, Rabrait-Lerman C, Poser BA, Ugurbil K, Boulant N. Magnetic field strength dependent SNR gain at the center of a spherical phantom and up to 11.7T. Magn Reson Med 2022;88(5):2131–2138. doi:10.1002/mrm.29391

18. Ocali O, Atalar E. Ultimate intrinsic signal-to-noise ratio in MRI. Magn Reson Med 1998;39(3):462–473. doi:10.1002/mrm.1910390317

19. Wiesinger F, Boesiger P, Pruessmann KP. Electrodynamics and ultimate SNR in parallel MR imaging. Magn Reson Med 2004;52(2):376–390. doi:10.1002/mrm.20183

20. Guerin B, Villena JF, Polimeridis AG, Adalsteinsson E, Daniel L, White JK, Wald LL. The ultimate signal-to-noise ratio in realistic body models. Magn Reson Med 2016. doi:10.1002/mrm.26564

21. Lee HH, Sodickson DK, Lattanzi R. An analytic expression for the ultimate intrinsic SNR in a uniform sphere. Magn Reson Med 2018;80(5):2256–2266. doi:10.1002/mrm.27207

22. Schnell W, Renz W, Vester M, Ermert H. Ultimate signal-to-noise-ratio of surface and body antennas for magnetic resonance imaging. IEEE Trans Antennas Propag 2000;48:418–428

23. Ohliger MA, Grant AK, Sodickson DK. Ultimate intrinsic signal-to-noise ratio for parallel MRI: electromagnetic field considerations. Magn Reson Med 2003;50(5):1018–1030. doi:10.1002/mrm.10597

24. Zhang B, Radder J, Giannakopoulos I, Grant A, Lagore R, Waks M, Tavaf N, Van de Moortele PF, Adriany G, Sadeghi-Tarakameh A, Eryaman Y, Lattanzi R, Ugurbil K. Performance of receive head arrays versus ultimate intrinsic SNR at 7 T and 10.5 T. Magn Reson Med 2024. doi:10.1002/mrm.30108

25. Ugurbil K, Auerbach E, Moeller S, Grant A, Wu X, Van de Moortele PF, Olman C, DelaBarre L, Schillak S, Radder J, Lagore R, Adriany G. Brain imaging with improved acceleration and SNR at 7 Tesla obtained with 64-channel receive array. Magn Reson Med 2019;82(1):495–509. doi:10.1002/mrm.27695

26. Feinberg DA, Beckett AJS, Vu AT, Stockmann J, Huber L, Ma S, Ahn S, Setsompop K, Cao X, Park S, Liu C, Wald LL, Polimeni JR, Mareyam A, Gruber B, Stirnberg R, Liao C, Yacoub E, Davids M, Bell P, Rummert E, Koehler M, Potthast A, Gonzalez-Insua I, Stocker S, Gunamony S, Dietz P. Next-generation MRI scanner designed for ultra-high-resolution human brain imaging at 7 Tesla. Nat Methods 2023. doi:10.1038/s41592-023-02068-7

27. Gruber B, Stockmann JP, Mareyam A, Keil B, Bilgic B, Chang Y, Kazemivalipour E, Beckett AJS, Vu AT, Feinberg DA, Wald LL. A 128-channel receive array for cortical brain imaging at 7 T. Magn Reson Med 2023. doi:10.1002/mrm.29798

28. Clement J, Gruetter R, Ipek O. A combined 32-channel receive-loops/8-channel transmit-dipoles coil array for whole-brain MR imaging at 7T. Magn Reson Med 2019;82(3):1229–1241. doi:10.1002/mrm.27808

29. Avdievich NI, Giapitzakis IA, Bause J, Shajan G, Scheffler K, Henning A. Double-row 18-loop transceive-32-loop receive tight-fit array provides for whole-brain coverage, high transmit performance, and SNR improvement near the brain center at 9.4T. Magn Reson Med 2019;81(5):3392–3405. doi:10.1002/mrm.27602

30. Avdievich NI, Nikulin AV, Ruhm L, Magill AW, Glang F, Henning A, Scheffler K. A 32-element loop/dipole hybrid array for human head imaging at 7 T. Magn Reson Med 2022;88(4):1912–1926. doi:10.1002/mrm.29347

31. Williams SN, McElhinney P, Gunamony S. Ultra-high field MRI: parallel-transmit arrays and RF pulse design. Phys Med Biol 2023;68(2). doi:10.1088/1361-6560/aca4b7

32. Wiggins GC, Polimeni JR, Potthast A, Schmitt M, Alagappan V, Wald LL. 96-Channel receive-only head coil for 3 Tesla: design optimization and evaluation. Magn Reson Med 2009;62(3):754–762. doi:10.1002/mrm.22028

33. Keil B, Blau JN, Biber S, Hoecht P, Tountcheva V, Setsompop K, Triantafyllou C, Wald LL. A 64-channel 3T array coil for accelerated brain MRI. Magn Reson Med 2013;70(1):248–258. doi:10.1002/mrm.24427

34. Adriany G, Auerbach EJ, Snyder CJ, Gozubuyuk A, Moeller S, Ritter J, Van de Moortele PF, Vaughan T, Ugurbil K. A 32-channel lattice transmission line array for parallel transmit and receive MRI at 7 tesla. Magn Reson Med 2010;63(6):1478–1485. doi:10.1002/mrm.22413

35. Lagore RL, Moeller S, Zimmermann J, DelaBarre L, Radder J, Grant A, Ugurbil K, Yacoub E, Harel N, Adriany G. An 8-dipole transceive and 24-loop receive array for non-human primate head imaging at 10.5 T. NMR Biomed 2021;34(4):e4472. doi:10.1002/nbm.4472

36. Shajan G, Hoffmann J, Budde J, Adriany G, Ugurbil K, Pohmann R. Design and evaluation of an RF front-end for 9.4 T human MRI. Magn Reson Med 2011;66(2):594–602. doi:10.1002/mrm.22808

37. Dubois M, Leroi L, Raolison Z, Abdeddaim R, Antonakakis T, de Rosny J, Vignaud A, Sabouroux P, Georget E, Larrat B, Tayeb G, Bonod N, Amadon A, Mauconduit F, Poupon C, Le Bihan D, Enoch S. Kerker Effect in Ultrahigh-Field Magnetic Resonance Imaging. Phys Rev X 2018;8:031083. doi:10.1103/PhysRevX.8.031083

38. Yan X, Gore JC, Grissom WA. Self-decoupled radiofrequency coils for magnetic resonance imaging. Nat Commun 2018;9(1):3481. doi:10.1038/s41467-018-05585-8

39. Lakshmanan K, Cloos M, Brown R, Lattanzi R, Sodickson DK, Wiggins GC. The "Loopole" Antenna: A Hybrid Coil Combining Loop and Electric Dipole Properties for Ultra-High-Field MRI. Concepts Magn Reson Part B Magn Reson Eng 2020;2020. doi:10.1155/2020/8886543

40. Raaijmakers AJ, Ipek O, Klomp DW, Possanzini C, Harvey PR, Lagendijk JJ, van den Berg CA. Design of a radiative surface coil array element at 7 T: the single-side adapted dipole antenna. Magn Reson Med 2011;66(5):1488–1497. doi:10.1002/mrm.22886

41. Wiggins G, Zhang B, Cloos M, Lattanzi R, Chen G, Lakshmanan K, Haemer G, Sodickson D. Mixing loops and electric dipole antennas for increased sensitivity at 7 Tesla.. Proceeding ISMRM, Salt Lake City, UT,USA 2013:2737

42. Pfrommer A, Henning A. The ultimate intrinsic signal-to-noise ratio of loop- and dipole-like current patterns in a realistic human head model. Magn Reson Med 2018;80(5):2122–2138. doi:10.1002/mrm.27169

43. Lattanzi R, Wiggins GC, Zhang B, Duan Q, Brown R, Sodickson DK. Approaching ultimate intrinsic signal-to-noise ratio with loop and dipole antennas. Magn Reson Med 2018;79(3):1789–1803. doi:10.1002/mrm.26803

44. Shajan G, Kozlov M, Hoffmann J, Turner R, Scheffler K, Pohmann R. A 16-channel dual-row transmit array in combination with a 31-element receive array for human brain imaging at 9.4 T. Magn Reson Med 2014;71(2):870–879. doi:10.1002/mrm.24726

45. Sadeghi-Tarakameh A, Radder J, Lagore R, Tavaf N, He X, Grant A, DelaBarre L, Vizioli L, Wu X, Auerbach E, Moeller S, Adriany G, Metzger G, Van de Moortele PF, Yacoub E, Ugurbil K, Eryaman Y. Safety Assessment of Custom-Built Multi-Channel RF Coils: EM Modeling Uncertainties. Proc Int Soc Mag Reson Med 2022;30:0583

46. International Electrotechnical C. International standard meI--. Particular requirements for the basic safety and essential performance of magnetic resonance equipment for medical diagnosis:. In: Commission IE, editor. Geneva, Switzerland; 2022.

47. Van de Moortele P-F, Ugurbil K. Very fast Multi Channel B1 calibration at high field in the small Flip angle regime.. Proc Int Soc Mag Reson Med (Annual meeting abstracts) 2009;17:367

48. Yarnykh VL. Actual flip-angle imaging in the pulsed steady state: a method for rapid three-dimensional mapping of the transmitted radiofrequency field. Magn Reson Med 2007;57(1):192–200. doi:10.1002/mrm.21120

49. Pruessmann KP, Weiger M, Scheidegger MB, Boesiger P. SENSE: sensitivity encoding for fast MRI. Magn Reson Med 1999;42(5):952–962

50. Barth M, Breuer F, Koopmans PJ, Norris DG, Poser BA. Simultaneous multislice (SMS) imaging techniques. Magn Reson Med 2016;75(1):63–81. doi:10.1002/mrm.25897

51. Moeller S, Auerbach E, van de Moortele P-F, Adriany G, Ugurbil K. fMRI with 16 fold reduction using multibanded multislice sampling. Proc Int Soc Mag Reson Med 2008;16:2366

52. Xu J, Moeller S, Auerbach EJ, Strupp J, Smith SM, Feinberg DA, Yacoub E, Ugurbil K. Evaluation of slice accelerations using multiband echo planar imaging at 3 T. Neuroimage 2013;83:991–1001. doi:10.1016/j.neuroimage.2013.07.055

53. Chen G, Collins CM, Sodickson DK, Wiggins GC. A method to assess the loss of a dipole antenna for ultra-high-field MRI. Magn Reson Med 2018;79(3):1773–1780. doi:10.1002/mrm.26777

54. Roemer PB, Edelstein WA, Hayes CE, Souza SP, Mueller OM. The NMR phased array. Magn Reson Med 1990;16(2):192–225

55. Le Garrec M, Gras V, Hang MF, Ferrand G, Luong M, Boulant N. Probabilistic analysis of the specific absorption rate intersubject variability safety factor in parallel transmission MRI. Magn Reson Med 2017;78(3):1217–1223. doi:10.1002/mrm.26468

56. Lattanzi R, Grant AK, Polimeni JR, Ohliger MA, Wiggins GC, Wald LL, Sodickson DK. Performance evaluation of a 32-element head array with respect to the ultimate intrinsic SNR. NMR Biomed 2010;23(2):142–151. doi:10.1002/nbm.1435

57. Ugurbil K, Xu J, Auerbach EJ, Moeller S, Vu AT, Duarte-Carvajalino JM, Lenglet C, Wu X, Schmitter S, Van de Moortele PF, Strupp J, Sapiro G, De Martino F, Wang D, Harel N, Garwood M, Chen L, Feinberg DA, Smith SM, Miller KL, Sotiropoulos SN, Jbabdi S, Andersson JL, Behrens TE, Glasser MF, Van Essen DC, Yacoub E, Consortium WU-MH. Pushing spatial and temporal resolution for functional and diffusion MRI in the Human Connectome Project. Neuroimage 2013;80:80–104. doi:10.1016/j.neuroimage.2013.05.012

58. Setsompop K, Gagoski BA, Polimeni JR, Witzel T, Wedeen VJ, Wald LL. Blipped-controlled aliasing in parallel imaging for simultaneous multislice echo planar imaging with reduced g-factor penalty. Magn Reson Med 2012;67(5):1210–1224. doi:10.1002/mrm.23097

59. Vizioli L, Moeller S, Dowdle L, Akcakaya M, De Martino F, Yacoub E, Ugurbil K. Lowering the thermal noise barrier in functional brain mapping with magnetic resonance imaging. Nat Commun 2021;12(1):5181. doi:10.1038/s41467-021-25431-8

60. Kozlov M, Turner R. Fast MRI coil analysis based on 3-D electromagnetic and RF circuit co-simulation. J Magn Reson 2009;200(1):147–152. doi:10.1016/j.jmr.2009.06.005

61. Lattanzi R, Sodickson DK. Ideal current patterns yielding optimal signal-to-noise ratio and specific absorption rate in magnetic resonance imaging: computational methods and physical insights. Magn Reson Med 2012;68(1):286–304. doi:10.1002/mrm.23198

62. Van de Moortele PF, Akgun C, Adriany G, Moeller S, Ritter J, Collins CM, Smith MB, Vaughan JT, Ugurbil K. B(1) destructive interferences and spatial phase patterns at 7 T with a head transceiver array coil. Magn Reson Med 2005;54(6):1503–1518. doi:10.1002/mrm.20708

63. Lagore RL, Grant A, DelaBarre L, Auerbach E, Waks M, Jungst S, Moeller S, Radder J, Sadeghi-Tarakameh A, Eryaman Y, Van de Moortele P-F, Adriany G, Ugurbil K. 128-channel brain imaging array with improved acceleration at 10.5 Tesla. Proc Int Soc Mag Reson Med 2023:1059

64. Sadeghi-Tarakameh A, Waks M, Grant A, Thotland J, Lagore RL, DelaBarre L, Auerbach E, Van de Moortele P-F, Adriany G, Ugurbil K, Eryaman Y. Boosting Central Head SNR at 10.5T: 32-channel Hybrid RF Coil Comprised of 25 Rx-only Loops and 7 TxRx NODES Dipoles. Proc Int Soc Mag Reson Med 2023:3913

65. Avdievich NI, Nikulin AV, Ruhm L, Magill AW, Henning A, Scheffler K. Double-row dipole/loop combined array for human whole brain imaging at 7 T. NMR Biomed 2022;35(10):e4773. doi:10.1002/nbm.4773

66. Vu AT, Auerbach E, Lenglet C, Moeller S, Sotiropoulos SN, Jbabdi S, Andersson J, Yacoub E, Ugurbil K. High resolution whole brain diffusion imaging at 7T for the Human Connectome Project. Neuroimage 2015;122:318–331. doi:10.1016/j.neuroimage.2015.08.004

67. Vaidya MV, Sodickson DK, Lattanzi R. Approaching Ultimate Intrinsic SNR in a Uniform Spherical Sample with Finite Arrays of Loop Coils. Concepts Magn Reson Part B Magn Reson Eng 2014;44(3):53–65. doi:10.1002/cmr.b.21268

68. Rooney WD, Johnson G, Li X, Cohen ER, Kim SG, Ugurbil K, Springer CS, Jr. Magnetic field and tissue dependencies of human brain longitudinal 1H2O relaxation in vivo. Magn Reson Med 2007;57(2):308–318. doi:10.1002/mrm.21122

## BIBLIOGRAPHY to SUPPORTING INFORMATION

1. Adriany G, Auerbach EJ, Snyder CJ, Gozubuyuk A, Moeller S, Ritter J, Van de Moortele PF, Vaughan T, Ugurbil K. A 32-channel lattice transmission line array for parallel transmit and receive MRI at 7 tesla. Magn Reson Med 2010;63(6):1478–1485.

2. Adriany G, Ritter J, Van de Moortele PF, Vaughan JT, Ugurbil K. Experimental verification of enhanced B1 Shim performance with a Z-encoding RF coil array at 7 tesla.". Proc Int Soc Mag Reson Med 2010;18:3831.

3. Kozlov M, Turner R. Analysis of RF transmit performance for a 7T dual row multichannel MRI loop array. Annu Int Conf IEEE Eng Med Biol Soc 2011;2011:547–553.

4. Shajan G, Kozlov M, Hoffmann J, Turner R, Scheffler K, Pohmann R. A 16-channel dual-row transmit array in combination with a 31-element receive array for human brain imaging at 9.4 T. Magn Reson Med 2014;71(2):870–879.

5. Feinberg DA, Beckett AJS, Vu AT, Stockmann J, Huber L, Ma S, Ahn S, Setsompop K, Cao X, Park S, Liu C, Wald LL, Polimeni JR, Mareyam A, Gruber B, Stirnberg R, Liao C, Yacoub E, Davids M, Bell P, Rummert E, Koehler M, Potthast A, Gonzalez-Insua I, Stocker S, Gunamony S, Dietz P. Next-generation MRI scanner designed for ultra-high-resolution human brain imaging at 7 Tesla. Nat Methods 2023.

6. Wu X, Tian J, Schmitter S, Vaughan JT, Ugurbil K, Van de Moortele PF. Distributing coil elements in three dimensions enhances parallel transmission multiband RF performance: A simulation study in the human brain at 7 Tesla. Magn Reson Med 2016;75(6):2464–2472.

7. Roemer PB, Edelstein WA, Hayes CE, Souza SP, Mueller OM. The NMR phased array. Magn Reson Med 1990;16(2):192–225.

8. Nelson JA, Stavis G. Impedance Matching, Transformers and Baluns. In: Reich HJ, editor. Very High-frequency Techniques. Volume 1; 1947. p 86–87.

9. Sadeghi-Tarakameh A, Radder J, Lagore R, Tavaf N, Adriany G, Ugurbil K, Eryaman Y. Safety assessment of custom-built multi-channel RF coils: EM modeling uncertainties. Proc Int Soc Mag Reson Med 2022:583.

10. Boulant N, Gras V, Amadon A, Le Bihan D. Workflow proposal for defining SAR safety margins in parallel transmission. 2018. p 295.

11. Steensma BR, Sadeghi-Tarakameh A, Meliado EF, van den Berg CAT, Klomp DWJ, Luijten PR, Metzger GJ, Eryaman Y, Raaijmakers AJE. Tier-based formalism for safety assessment of custom-built radio-frequency transmit coils. NMR in biomedicine 2023;36(5):e4874.

12. Le Garrec M, Gras V, Hang MF, Ferrand G, Luong M, Boulant N. Probabilistic analysis of the specific absorption rate intersubject variability safety factor in parallel transmission MRI. Magn Reson Med 2017;78(3):1217–1223.

13. Sadeghi-Tarakameh A, DelaBarre L, Lagore RL, Torrado-Carvajal A, Wu X, Grant A, Adriany G, Metzger GJ, Van de Moortele PF, Ugurbil K, Atalar E, Eryaman Y. In vivo human head MRI at 10.5T: A radiofrequency safety study and preliminary imaging results. Magn Reson Med 2020;84(1):484–496.

14. Graesslin I, Homann H, Biederer S, Bornert P, Nehrke K, Vernickel P, Mens G, Harvey P, Katscher U. A specific absorption rate prediction concept for parallel transmission MR. Magn Reson Med 2012;68(5):1664–1674.

15. Eichfelder G, Gebhardt M. Local specific absorption rate control for parallel transmission by virtual observation points. Magn Reson Med 2011;66(5):1468–1476.

